# A shared theta-rhythmic process for selective sampling of environmental information and internally stored information

**DOI:** 10.1101/2024.11.26.625454

**Authors:** Paul J. Cavanah, Ian C. Fiebelkorn

## Abstract

Selective attention is the collection of mechanisms through which the brain preferentially processes behaviorally important information. Many everyday tasks, such as shopping for groceries, require selective sampling (i.e., attention-related sampling) of both external information (i.e., information from the environment) and internally stored information (i.e., information being maintained in working memory). While there is clear evidence that selective sampling of external information is influenced by internally stored information (and vice versa), the extent to which selective sampling of external and internal information share neural resources remains a focus of debate. Previous research has linked theta-rhythmic (3–8 Hz) neural activity in higher-order (e.g., frontal cortices) and sensory regions to theta-rhythmic changes in behavioral performance during selective sampling. Here, we used EEG and a dual-task design (i.e., a task that required both external and internal information), in male and female humans, to directly compare theta-dependent fluctuations in behavioral performance during external sampling with those during internal sampling. Our findings are consistent with a shared theta-rhythmic process for selectively sampling external information or internal information. This theta-rhythmic sampling is associated with both phase-dependent changes in sensory responses (i.e., as measured with the N1 component) and phase-dependent changes in interactions between external and internal information. The theta phase associated with weaker sensory responses and relatively worse behavioral performance (i.e., the ‘bad’ phase) is also associated with a slowed perceptual decision-making process (as measured with the CPP component), specifically during dual-task trials when to-be-detected external information matches to-be-remembered internal information.

**SIGNIFICANCE STATEMENT:** Most everyday tasks require information from both the external environment and internal memory stores; however, the extent to which selective processing of external and internal information rely on shared neural mechanisms and resources remains a subject of debate. Recent work has demonstrated attention-related, theta-rhythmic fluctuations (3–8 Hz) in neural activity and behavioral performance, perhaps reflecting the temporal coordination of competing functions (e.g., attention-related sampling and shifting). Here, we used EEG and a dual-task design to provide evidence of a shared, theta-rhythmic process for alternately boosting the sampling of either external or internal information. This shared, theta-rhythmic process also modulates interactions between external and internal information on dual-task trials, when these sources of information compete for limited processing resources.

## INTRODUCTION

Given limited processing resources, the brain uses a collection of filtering mechanisms to preferentially process behaviorally important information. This essential cognitive function, broadly referred to as selective attention, is often investigated in the context of sampling information from the environment (i.e., external information)^1, 2^; however, everyday tasks, such as shopping for groceries, also require the selective sampling of internally stored information. For example, we can selectively sample behaviorally important information that is being maintained in working memory^3, 4^. This internal sampling process has been characterized as internal selective attention^5–11^. While selective sampling of external information and selective sampling of internal information are often investigated separately, there is increasing interest in the relationship between them^5, 6, 8, 12–15^. Previous research has demonstrated interactions between external and internal sampling, often by utilizing dual-task designs that potentially create competition between these processes^10, 16–22^. For example, there can be a relative advantage for an external visual target that matches a to-be-remembered item being concurrently maintained in working memory^19, 20, 23, 24^. On the other hand, increasing working memory load (e.g., the number of to-be-remembered items) can decrease the detection of near-threshold visual targets^25–27^. While such findings are consistent with shared neural resources^28–31^, the extent to which selective sampling of external information and selective sampling of internal information share neural resources remains a focus of debate^29, 32–39^.

Recent evidence suggests that theta-rhythmic (3–8 Hz) neural activity shapes both external sampling of environmental information and internal sampling of information being maintained in working memory^40–56^. In the context of external sampling, preferential processing (or sampling) at specific locations in space (i.e., spatial attention) is sometimes likened to a spotlight that scans the visual environment, pausing to illuminate behaviorally important, external information. Whereas the classic view of this spotlight is that it can be deployed continuously, newer findings indicate that it dims about 3–8 times per second^40–50^. That is, attention-related changes in both neural activity and behavioral performance fluctuate at a theta frequency. Similar to these findings, neural and behavioral measures associated with to-be-remembered items—during working memory tasks—also fluctuate at a theta frequency^51–57^. Such theta-rhythmic fluctuations might reflect the temporal coordination of shared neural resources, isolating neural activity associated with different functions (or representations) over time^40^. For example, we and others have proposed that theta-rhythmic neural activity coordinates sampling (i.e., sensory functions) and shifting (i.e., motor functions) within brain regions that direct both selective attention and goal-directed orienting movements (i.e., within the ‘Attention Network’)^40, 41, 58^.

Here, we used electroencephalography (EEG) and a dual-task design (i.e., a behavioral task that requires both external and internal sampling) to investigate whether there is a shared, theta-rhythmic process for selectively boosting the sampling of either external or internal information. We specifically compared the relationship between the phase of theta-band activity and behavioral performance across trials that probed external sampling (i.e., with a noise-added visual target) and trials that probed internal sampling (i.e., by retro-cueing a to-be-remembered item). That is, we tested whether selective enhancement of external information and selective enhancement of internal information occur at either the same phase or different phases of theta-rhythmic neural activity. Figure 1 illustrates different ways in which external and internal sampling might be phasically coordinated during a shared, theta-rhythmic sampling process. We further tested whether theta-rhythmic sampling influences behavioral and electrophysiological markers of interactions between internal and external information. For example, behavioral and electrophysiological interactions that occur when to-be-detected external information matches to-be-remembered internal information (i.e., so-called ‘match effects’)^16, 17^. Our findings are consistent with a shared theta-rhythmic process for selectively sampling environmental information or internally stored information. This theta-rhythmic sampling process is associated with (i) phase-dependent changes in neural activity attributable to sensory processing (as measured with the N1 component) and (ii) phase-dependent changes in neural activity attributable to decision making (as measured with the CPP component). Effects on decision making specifically occur during trials that require both external and internal sampling (i.e., during dual-task trials).

**Figure 1.**
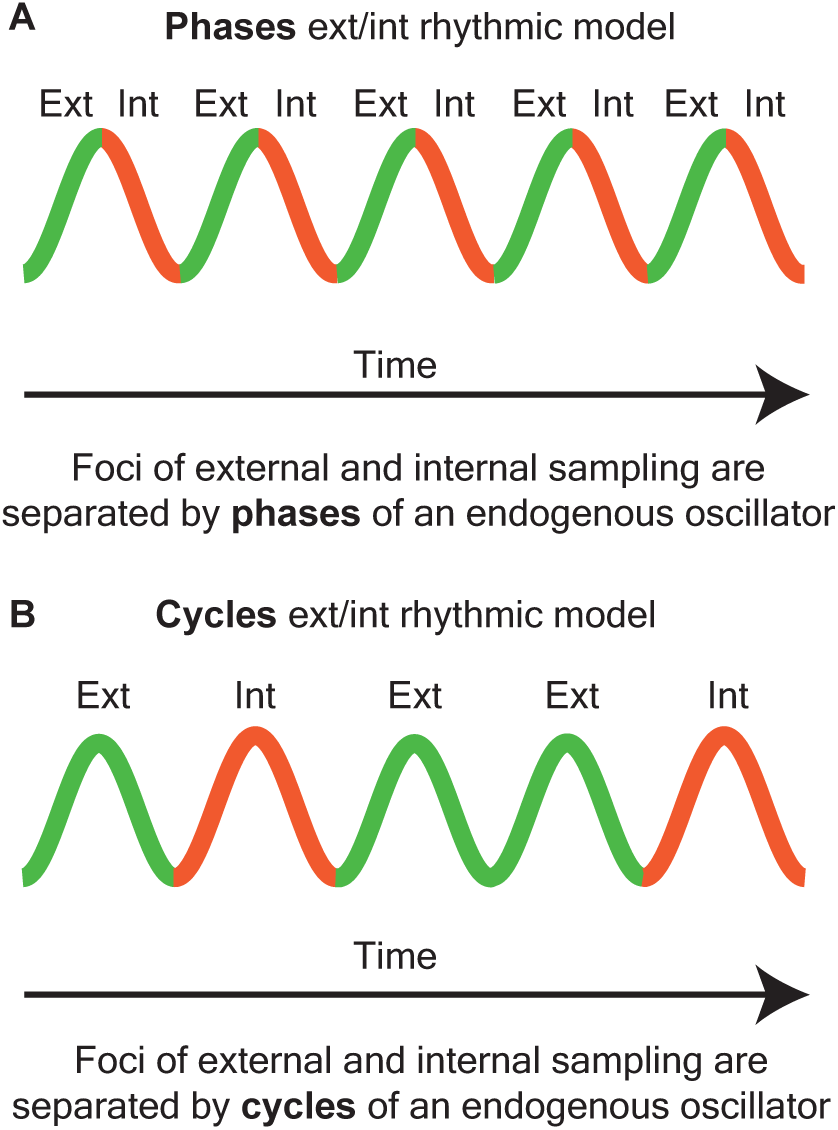
Suggested models for the coordination of external and internal sampling via a shared endogenous rhythm. (A) Phases model: Within a single cycle of an endogenous oscillation, the sampling of external or internal information is separated by the phase of the cycle. (B) Cycles model: Single cycles are dedicated to processing a single focus at a time (either external or internal), with inter-cycle periods (i.e., troughs of sampling) serving as opportunities to shift between foci. The “cycles” model predicts that there is the same peak phase of sampling for both external and internal information, while the “phases” model predicts that there are different peak phases of sampling for external and internal information.

## MATERIALS AND METHODS

### Participants

Thirty-one individuals (18 females), aged 19-33 years old (average 23.5), participated in the experiment. Participants completed two sessions of approximately two hours each, separated by at least 24 hours and no more than two weeks. For their time, the participants received $15 per hour. All participants had normal or corrected-to-normal vision and no history of neurological conditions. In accordance with the Declaration of Helsinki, each participant gave written informed consent prior to data collection. The study protocol was approved by the University of Rochester Research Participants Review Board. We excluded six participants from the analyses because of excessive noise in their EEG signals, resulting in the rejection of greater than 20 percent of trials (see below for a description of preprocessing).

### Behavioral task

The behavioral task is illustrated in Figure 2A. Participants were seated in a quiet, light-attenuated recording booth in front of a 24-inch LCD monitor (ASUS Predator, 240 Hz refresh rate). Head position and viewing distance were controlled with a fixed chin rest, positioned 57 cm from the screen. At the start of each session, the experimenter explained the behavioral task to the participant, who was then given practice trials until they demonstrated an understanding of the task. The participants used a keyboard to initiate trials (i.e., trials were self-paced) and to give responses. Throughout all trial stages, until a response was given, participants were required to maintain fixation on a small, gray square (0.5 x 0.5 degrees) at the center of the screen. We measured eye position with an EyeLink 1000 Plus (SR Research, Ontario, Canada), which samples binocularly at 2000 Hz. If the position of a participant’s eyes deviated by more than one degree from central fixation, the trial was aborted and a new trial was initiated. Aborted trials were replaced, such that each participant completed the same number of trials.

**Figure 2.**
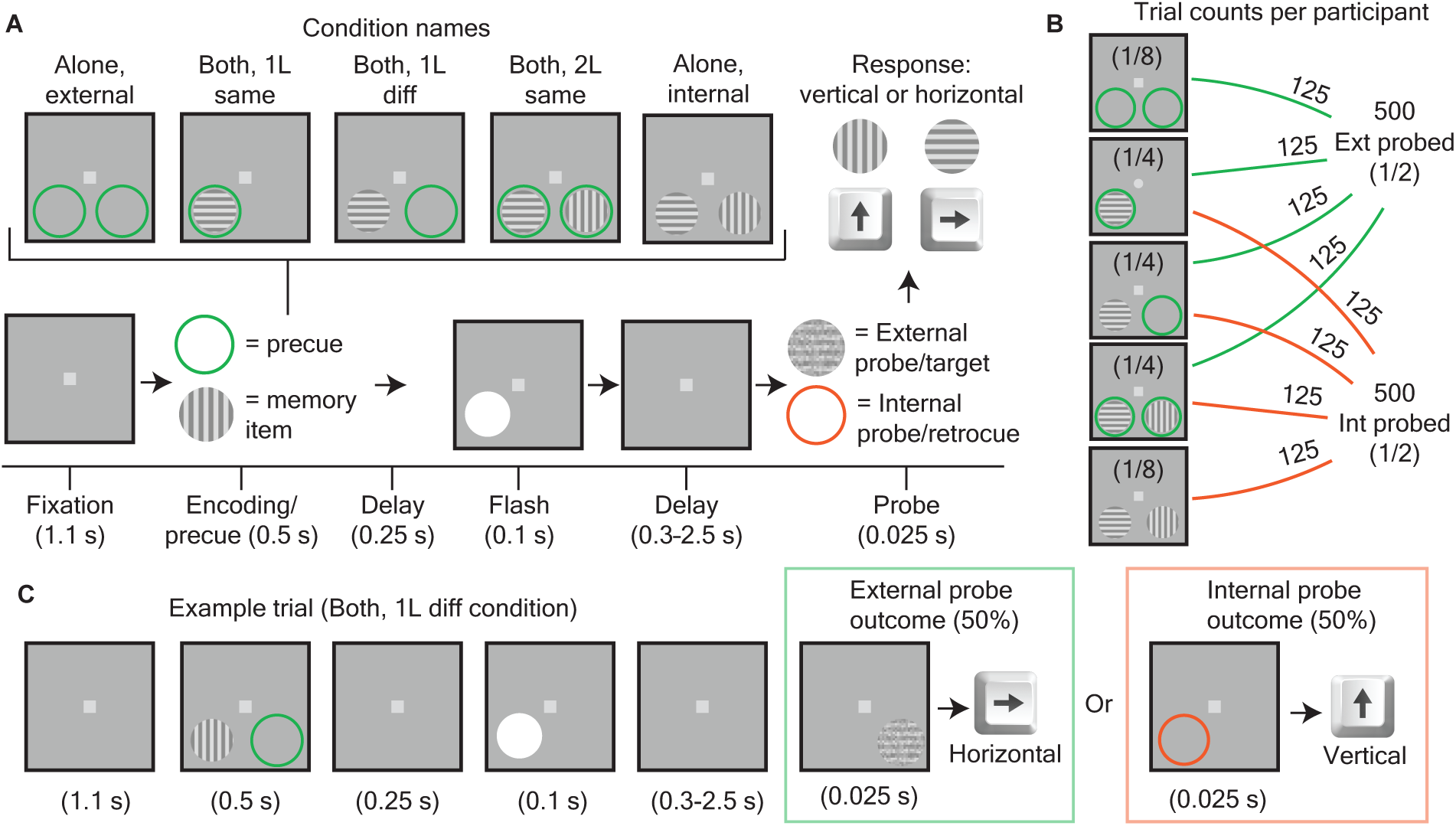
External/internal dual task. (A) Task schematic. Each trial began with a 1.1s fixation period, followed by a 0.5s period in which one of five encoding/precue arrays could be displayed (either external alone, internal alone, or a both condition). The green circles at this stage were pre-cues for possible external probes (low-contrast gratings) later in the trial. The gratings at this stage were memory items, which could be later tested by internal probes (retro-cues). After the encoding/pre-cue period, a short, fixed delay (0.25s), a flash event (0.1s), and then a variable delay (0.3-2.5s) occurred. At the probe stage (0.025s), either the external or internal modality could be probed. In ‘both’ trials, the probe types were equiprobable. The response to external and internal probes was indicated in the same way—either an “up arrow” (vertical orientation) report or a “right arrow” (horizontal orientation) report. (B) Trial counts per condition. For each encoding/cueing condition, the number of trials are shown. There were 1000 total trials per participant, with eight behaviorally distinct conditions (four external and four internal), equaling 125 trials per condition, and an even split of 500 trials between external and internal probes. (C) An example trial, specifically for the ‘both, 1L diff’ condition. At one location, the participant can expect an external probe with 50% probability, and at the other location, the participant can expect an internal probe with 50% probability. Trials ended with either an external probe or an internal probe.

We used Presentation software (Neurobehavioral Systems, Albany, CA, USA) to control the presentation of stimuli and to log responses. Trials consisted of seven sequential stages (see Fig. 2A): a 1.1s fixation/baseline period, a 0.5s cueing/encoding period, a 0.25s delay, a 0.1s flash event, a variable delay of 0.3–2.5s (sampled uniformly), a 0.025s probe, and a response period (with a response cutoff at 2.0s). All task stimuli were presented at either of two locations: one location to the right of central fixation and one location to the left of central fixation. Both locations were below the horizontal meridian, at a distance of 8 degrees from central fixation. Gratings (i.e., memory items and external probes) were 4 degrees in diameter and cues (i.e., pre-cues and retro-cues) were 4.4 degrees in diameter. During the encoding/pre-cueing period, combinations of two types of stimuli could be presented: green circles (i.e., pre-cues to sample external information) and/or square-wave gratings (i.e., to-be-remembered items). Green circles during the encoding/pre-cueing period indicated the potential location(s) for an upcoming, noise-added grating, which served as a visual target (i.e., an external probe). Noiseless gratings presented during the encoding/pre-cueing period were memory items to be maintained in anticipation of a potentially upcoming orange circle, which served as a retro-cue^59^ (i.e., an internal probe). The encoding period was followed by a brief, 0.25s delay and then a task-irrelevant, 0.1s flash event (i.e., a white circle). This flash event occurred at a previous stimulus location, with equal probability when there were two stimulus locations. Flash events, which have been similarly utilized by previous studies, create consistent sampling patterns across trials when there are multiple target locations^42, 45, 51^. After the flash event, there was a variable delay, 0.3–2.5s, when participants needed to (i) maintain neural representations of to-be-remembered gratings (i.e., until a retro-cue), (ii) boost sensory processing in anticipation of an upcoming, noise-added grating (i.e., a visual target), or (iii) both. Finally, the variable delay was followed by either a target (i.e., a noise-added, vertical or horizontal grating) or a retro-cue^59^ (i.e., an orange circle). The response scheme was the same for both externally probed trials (i.e., target trials) and internally probed trials (i.e., retro-cue trials): participants made a two-alternative forced choice, by pressing the ‘right arrow’ on the keyboard to indicate a horizontal grating and the ‘up arrow’ on the keyboard to indicate a vertical grating. On dual-task trials, which required both external and internal sampling (i.e., ‘both’ trials), the probe type was determined randomly, with equal probability. That is, while participants anticipated the possibility of either an external probe (i.e., a visual target) or an internal probe (i.e., a retro-cue) on dual-task trials, only one or the other was presented following the variable delay. On trials where only external or internal information was presented during the encoding/pre-cueing period (i.e., ‘alone, external’ and ‘alone, internal’ trials), the probe type was always the same as the information (e.g., memory items on an ‘alone, internal’ trial were always followed by a retro-cue). The behavioral task included the following trial types (see Fig. 2A): external sampling only (i.e., ‘alone, external’), internal sampling only (i.e., ‘alone, internal’), both external and internal sampling at one location (i.e., ‘both, 1L same), both external and internal sampling at different locations (i.e., ‘both, 1L different’), and both external and internal sampling at both locations (‘both, 2L same’).

The deployment of attentional resources during external sampling was promoted by using a noise-added grating as the visual target. To generate noise-added gratings, a matrix of uniformly distributed random values was added to square-wave gratings. Importantly, this matrix was multiplied by a scalar ‘difficulty constant,’ with a value of zero meaning no noise. This scalar determined the contrast of the dark/light stripes and served as the difficulty parameter for our staircasing procedure. The noise-added gratings (or noiseless gratings, in the case of to-be-remembered items) were min-max normalized to grayscale RGB values between 140 and 220. Every grating stimulus presented during the experiment therefore had the same mean grayscale RGB value as the background (i.e., 180), and the same minimum (i.e., 140) and maximum (i.e., 220) values. This ensured that the brightness of the gratings was always the same, differing only in the coherence of the vertical or horizontal stripes. Figure 2C shows an example of a noise-added grating (i.e., an external probe/visual target). The amount of noise added to the external probe was calibrated for each participant to achieve 80% accuracy across ‘alone, external’ trials. Here, we used a staircasing algorithm: on every trial, we calculated the accuracy (i.e., the proportion of correct trials) of all previous trials in the ‘alone, external’ condition (throughout the session). If the accuracy was lower than 0.78, the ‘difficulty constant’ (see above) was lowered. If the accuracy was higher than 0.82, the ‘difficulty constant’ was increased. To maximize the effectiveness of this procedure, we started each experiment with a ‘baseline’ block of 50 ‘alone, external’ trials (and no other trial types). Data from this baseline established a starting level of difficulty but were excluded from all analyses.

### Behavioral analyses

We first compared accuracy and response times between cue/memory conditions and delay lengths (Fig. 3). For all analyses, we split dual-task trials depending on whether external or internal sampling was probed. In accordance with previous research that suggests that interactions between external and internal sampling change over time, we also split trials depending on median delay length (1.4s)^16^. To determine significant behavioral differences, we conducted repeated measures ANOVAs with follow-up t-tests. The significance threshold for all statistical testing was p < 0.05. In Figure 3 and Supplementary Figure 1, we tested the behavioral difference between cue/memory conditions (‘alone’, ‘1L same’, ‘1L diff’, and ‘2L same’) and ‘short’ (less than median delay length) versus ‘long’ (longer than median delay length) trials (i.e., a 4×2 repeated measures, two-way ANOVA) for each probe (external and internal) and behavioral (accuracy and response time) type. In Figure 4, we further compared behavioral performance for (i) ‘both’ trials when the external probe (i.e., a noise-added grating) was a ‘match’ for a previously presented memory item (i.e., a noiseless grating), and relative to (ii) ‘both’ trials when the external probe was a ‘nonmatch’ for a previously presented memory item. Depending on the specific condition, matches in orientation could occur either at the same spatial location or at different spatial locations (see Fig. 4A). Trials were further binned based on whether the delay period duration was greater than or less than the median (i.e., 1.4s). In Figure 4, we tested the behavioral difference for the three ‘both’ conditions across ‘match’ versus ‘nonmatch’ and ‘short’ versus ‘long’ trials (i.e., a 3×2×2 repeated measures, three-way ANOVA). Finally, to more precisely visualize temporal differences in behavioral performance between ‘alone’ versus ‘both’ (Fig. 3C, 3F, S1C, S1F) and ‘match’ and ‘non-match’ (Fig. 4C) conditions over time, we calculated accuracy and response time within 0.6s bins with a step size of 0.01s (e.g., 0.3–0.9s, 0.31–0.91s, 0.32–0.92s, etc…). For alone vs. both windowed analyses (Fig. 3C, F, I, L), we combine ‘both’ conditions (i.e., ‘1L same’, ‘1L diff’, ‘2L same’). For match versus nonmatch windowed analyses (Fig. 4C), we combined the same-location ‘both’ conditions (i.e., the conditions that showed significant ‘match’ effects in the previous analysis, see Fig. 4B). For these windowed accuracy comparisons, we used a cluster-based approach^60^ to control for multiple comparisons between windows (see “cluster-based statistics in phase-behavior, windowed behavior, and ERPs” section).

**Figure 3.**
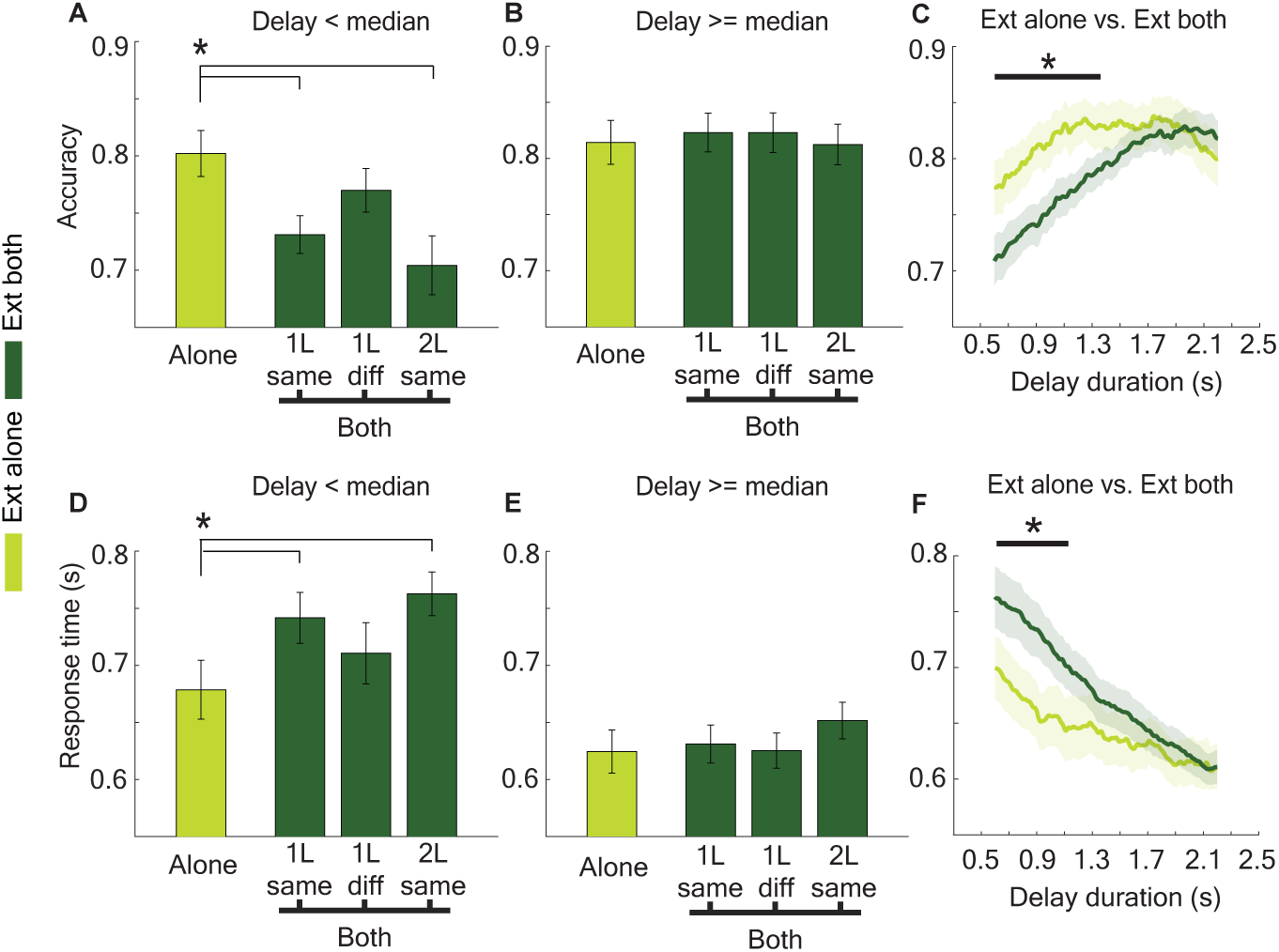
Internal sampling interferes with perceptual sampling at short delays. (A) Accuracy for external alone and external both (“1L same”, “1L diff”, and “2L”) trials in which the delay duration was less than the median (< 1.4s). (B) Same as A, but for trial delays longer than the median (> 1.4s). (C) Delay-duration windowed accuracy for external alone and external both (the ‘both’ conditions have been combined). Window length was 0.6s, window step was 0.01s. (D-F) Same as (A-C), but for response times (RTs). Asterisks between bars indicate p < 0.05 from a two-tailed t-test comparison. Solid lines indicate p < 0.05 clusters of significant values for the difference between the depicted conditions.

**Figure 4.**
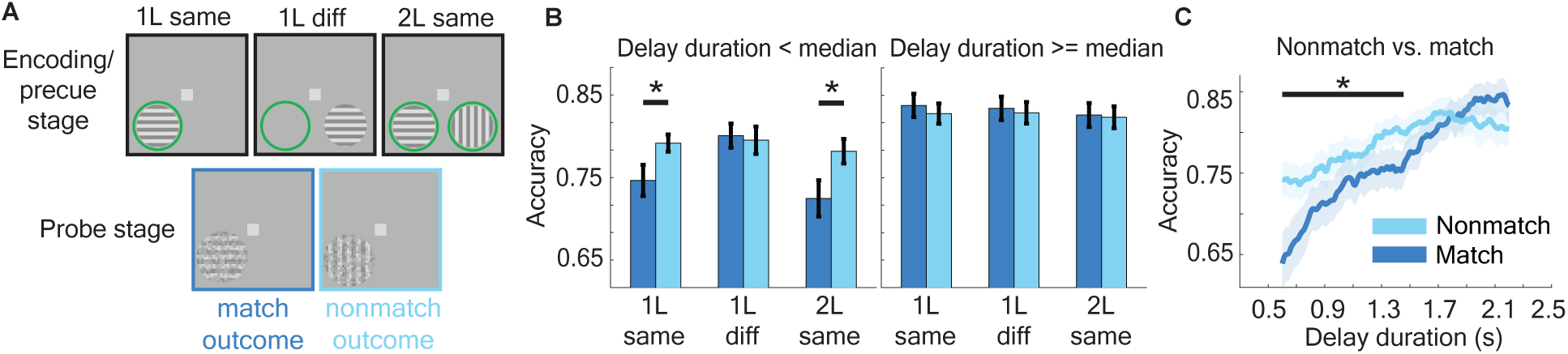
When the memory item matches the external probe at short delays, there is significantly lowered discriminability of the external probe. (A) Depiction of the ‘both’ conditions and their match/nonmatch outcomes. When the orientation of the to-be-remembered item (i.e., the previously presented memory item) at a particular location matches the orientation of the external probe, this is considered a ‘match.’ Importantly, the ‘1L diff’ condition doesn’t have a memory item at the same location as the external probe, so the match status is determined by a memory item presented at the opposite location as the external probe. (B) Accuracies in each condition. There was a significant decrease in accuracy for trials in which the delay was shorter than the median (< 1.4s) and the memory item previously presented at the same location matched the external probe. Significance was determined by a two-tailed t-test (p < 0.05). (C) Windowed accuracy for same-location match and nonmatch trials. Window length was 0.6s, window step was 0.01s. Solid lines indicate p < 0.05 clusters of significant values for the difference between the depicted conditions.

### Data acquisition and preprocessing

We recorded electroencephalographic (EEG) signals at a rate of 2048 Hz, using a 128-channel ActiveTwo BioSemi system (Amsterdam, the Netherlands). For all preprocessing and data analyses, we used a combination of customized MATLAB functions (MathWorks, Natick, MA, USA) and the Fieldtrip toolbox (Donders Institute for Brain, Cognition, and Behavior, Radboud University Nijmegen, the Netherlands)^61^. We downsampled the EEG data to 512 Hz and re-referenced using the 128-electrode average (i.e., an average reference). We then used a discrete Fourier Transform (DFT) filter—applied at 60, 120, and 180 Hz—to remove line noise. After epoching trials from 4.5 seconds before the probe to 1.0 second after the probe, we linearly detrended and demeaned the trial-level data. Trials with either eye blinks or saccades (> 1 degree) were aborted during recording sessions, so there was no need for rejection/correction based on eye artifacts during the initial preprocessing. We used voltage threshold of ±100 μV for identifying trials/channels with other noise transients, interpolating electrodes that exceeded this threshold using the nearest neighbor spline^62^. If more than 10% of electrodes needed to be interpolated on a single trial, we excluded the trial from further analyses^10^. If more than 20% of trials were excluded for a given participant, we excluded all the participant’s data from further analyses (n = 6). The remaining participants (n = 25) had an average of 6% of trials removed during artifact rejection, leaving an average of 939 trials per participant (across all conditions).

### Measuring phase-behavior relationships

We calculated phase-behavior relationships to determine if RTs and/or accuracy varied as a function of frequency-specific phase, measured just prior to probe onset (i.e., during the variable delay, just prior to the presentation of either a visual target or a retro-cue)^63^. Here, we first used Morlet wavelet convolution to derive frequency-specific phase measurements for each trial (from 3–55 Hz). To avoid contamination of the phase estimates from either flash-evoked visual responses or probe-evoked visual responses, (i) trials with delay periods less than 0.75s were excluded from the phase-behavior analyses (i.e., to avoid probe-evoked visual responses) and (ii) the time point for each phase measurement (i.e., the center of the wavelet) was half a wavelet width (i.e., half the temporal extent of the wavelet) from probe onset. That is, the wavelet was fit for each frequency such that the last time point included for calculation of the phase measurement was the time point just prior to probe onset. For example, phase for the 4 Hz, 2-cycle wavelet—with a temporal extent of 0.5s—was measured 0.25s prior to probe onset (i.e., the wavelet was centered 0.25s prior to probe onset). We calculated phase estimates from 3–8 Hz, in 1-Hz increments, and from 9–55 Hz, in 2-Hz increments. The number of cycles per wavelet was 2 for wavelets from 3–8 Hz and increased logarithmically from 2 to 5 cycles for wavelets from 9–55 Hz.

After obtaining electrode- and frequency-specific phase measurements (from 3–55 Hz), we paired trial-specific phase measurements with trial-level RTs. Normalization of RTs was done within participant and condition of interest. We then binned z-RTs by electrode- and frequency-specific phase, using bins with a width of 180 degrees (e.g., 0 to 180 degrees), shifted in 10-degree steps (e.g., 0 to 180 degrees, 10 to 190, etc.). Following this binning procedure, we calculated the median RT for each bin, providing z-RT as a function of frequency-specific phase for each electrode and condition of interest. We next averaged the participant-level phase-RT functions (n = 25) to determine the frequencies and electrodes with the strongest phase-RT relationships. We also calculated phase-accuracy functions (Fig. S3) using a nearly identical procedure, except that the proportion of correct trials was calculated for each phase bin, rather than median z-RT. Because accuracy was at ceiling for internal-probed trials (i.e., retro-cued trials), we only calculated phase-accuracy functions for external-probed trials (i.e., target trials). Similar to previous studies^44^ we hypothesized that the phasic modulation of behavioral performance would have a “peak” and a “trough” (i.e., a “good” phase and a “bad” phase) that were separated by approximately 180 degrees. The strength of the grand-averaged phase-behavior functions was therefore summarized by the amplitude of a one-cycle sine wave. We specifically used a discrete Fourier transform (DFT) applied to each grand-averaged function (at each frequency and electrode), with the absolute value of the second component of the DFT output being used to approximate the amplitude of a one-cycle sine fit to the phase-behavior function^44, 51, 63, 64^.

To determine the statistical significance of phase-behavior relationships, we randomly shuffled the observed phases and RTs (1500 permutations) before recalculating the strength of the phase-behavior relationships (following the same procedure outlined above). We then compared the observed values to the null distribution of randomized values, separately for each condition, to calculate p-values (i.e., the proportion of values in the null distribution that exceeded the observed value) for all electrodes and frequencies. Finally, we applied cluster-correction to correct for multiple comparisons, which has been widely used for EEG and MEG analyses^60^ (see “cluster-based statistics in phase-behavior, windowed behavior, and ERPs” section).

To determine whether the “peak” phases (i.e., the phases associated with better behavioral performance) differed between external and internal sampling, we compared the distributions of “peak” phases across participants for the conditions of interest (i.e., external sampling vs. internal sampling). Here, the analyses were limited to the frequencies and electrodes where the strongest phase-behavior relationships were observed during previous analyses. We used a circular Watson-Williams test to determine whether the participant-level distribution of “peak” phases during external sampling significantly differed from the participant-level distribution of “peak” phases during internal sampling (Fig. 5E and Fig. 6C)^51^.

**Figure 5.**
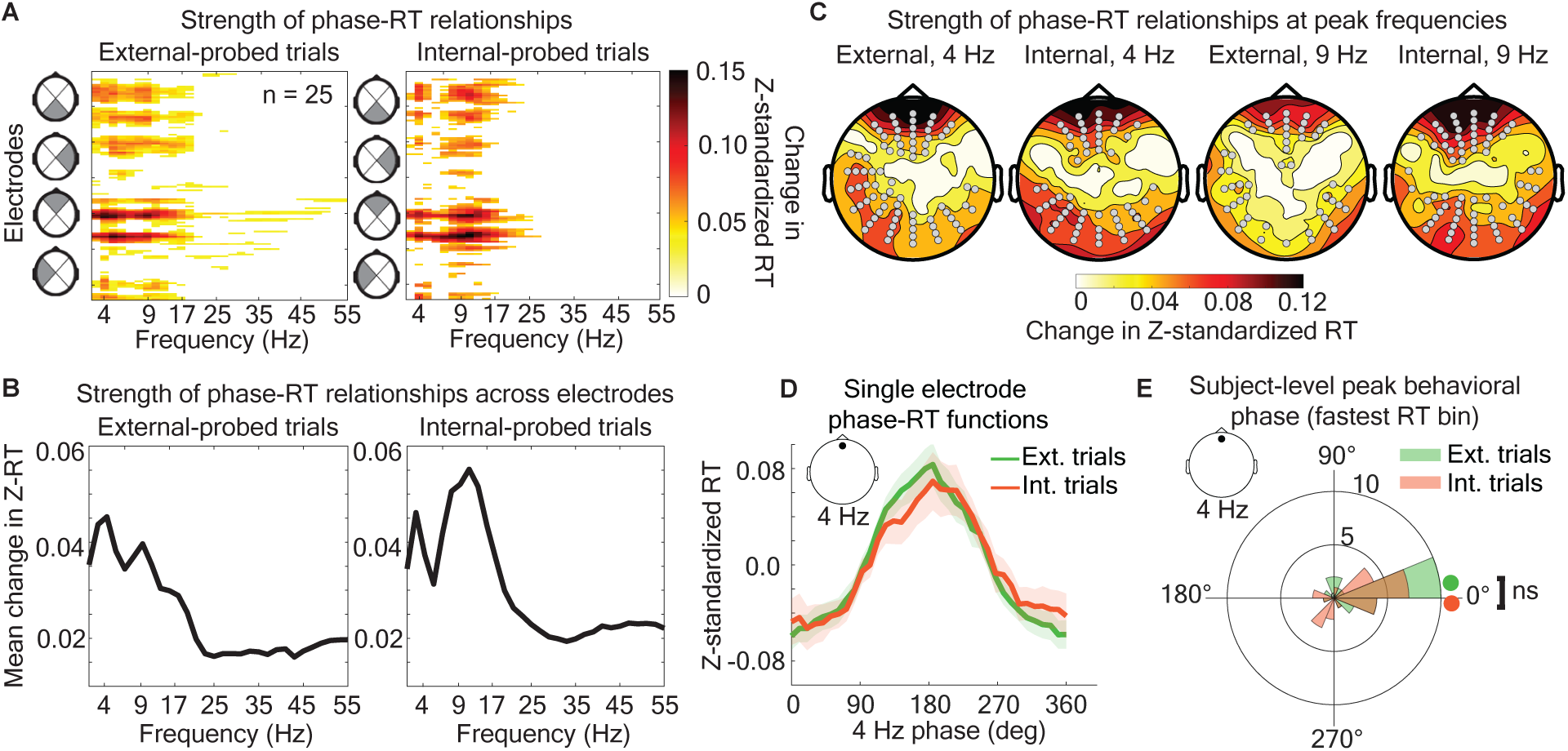
The pre-probe EEG phase similarly predicts external and internal performance. (A) Strength of phase-RT relationships are shown for a range of frequencies (from 3 to 55 Hz) and electrodes (128 total). All values that are not part of p<0.05 significant clusters are zeroed. *Left*: Strength of phase-RT relationships in response to external probes (low-contrast gratings). *Right:* Strength of phase-RT relationships in response to internal probes (retro-cues). (B) Phase-RT strength averaged across all electrodes for each frequency, separately for external-(left) and internal-probe (right) trials. Phase-RT strength, when averaging across electrodes, peaked at 4 Hz and 9 Hz. (C) Topographies of phase-RT strength at 4 Hz and 9 Hz. (D) Grand-averaged phase-RT functions for external-(green) and internal-probe (orange) trials from a single electrode. Inset depicts electrode location (frontal) and frequency (4 Hz). (E) The phase at which behavioral performance was the best for each participant, separately for external-(green) and internal-probed (orange) trials. Colored circles depict the grand-averaged, peak phases (i.e., represent the peak phases of the data shown in D). The radial axis depicts the number of participants in each peak-phase bin. Inset depicts electrode location (frontal) and frequency (4 Hz).

**Figure 6.**
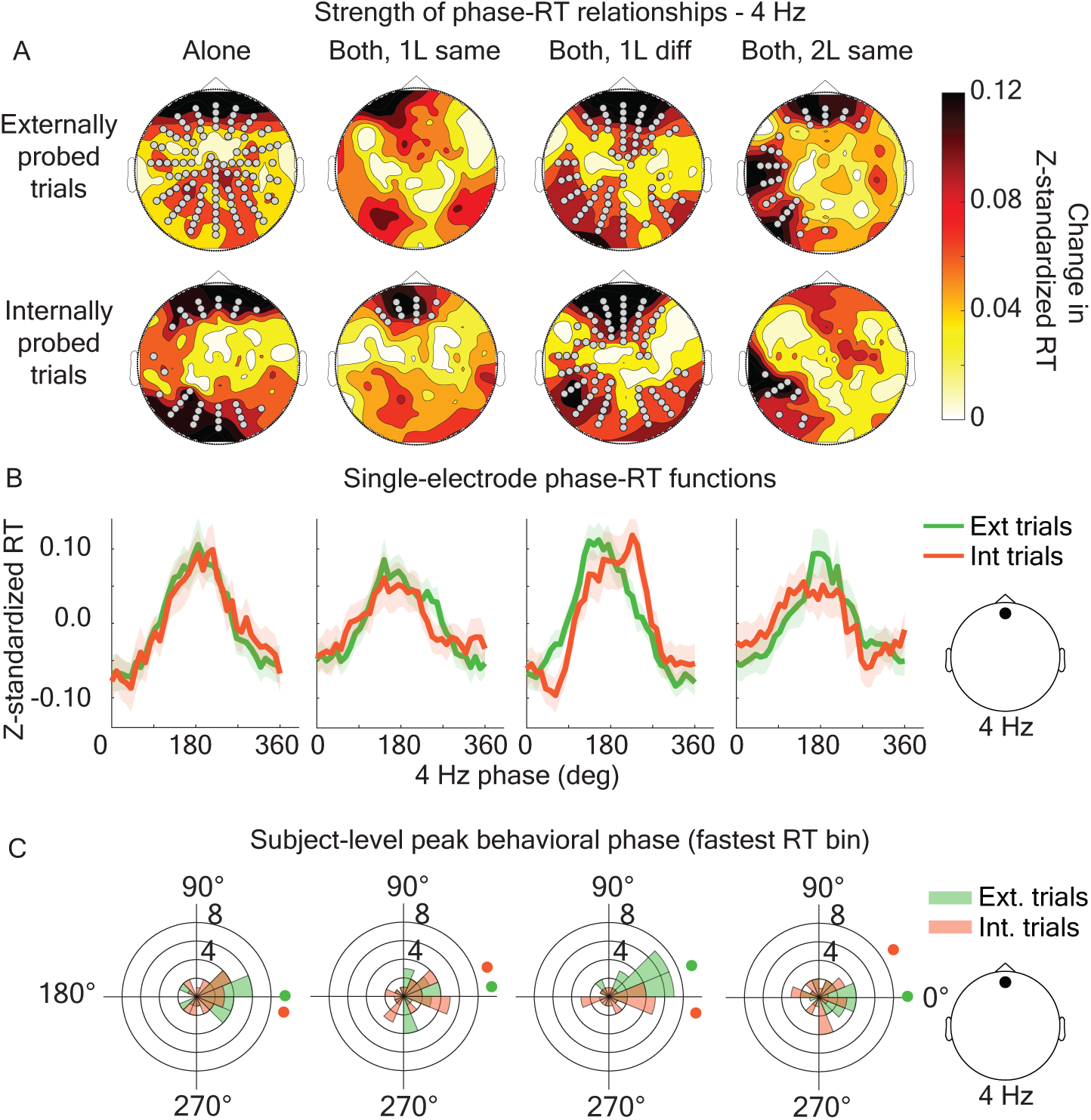
Phase-RT relationships are consistent across conditions. (A) Topographies and significant electrodes for phase-RT relationships across the different ‘alone’ and ‘both’ (i.e., dual-task) conditions, separately for external- and internal-probe trials. (B) Phase-RT functions for external-(green) and internal-probe (orange) trials, for the same conditions as (A). Inset on far right depicts electrode location (frontal) and frequency (4 Hz) of the phase measurement. (C) The phase at which behavioral performance was the best for each participant for the same conditions as (A) and (B). Solid colored circles depict the grand-averaged, peak phase (represent the peak phases of the data shown in B). The radial axis depicts the number of participants in each peak-phase bin. Inset on far right depicts electrode location (frontal) and frequency (4 Hz).

### Excluding trials with microsaccades from phase-behavior relationships

Like behavioral performance, previous research has demonstrated that the likelihood of microsaccades (i.e., fixational eye movements) fluctuates as a function of theta phase^65^. Previous work has also shown that phase-behavior relationships persist after excluding trials with microsaccades, indicating that phase-behavior relationships are not attributable to microsaccades^44^. We conducted a control analysis with the present data to similarly confirm that phase-behavior relationships are not attributable to microsaccades. That is, we re-calculated phase-behavior relationships after excluding trials with microsaccades during the delay period (i.e., in the pre-probe period).

To distinguish microsaccadic events, we set the EyeLink online parser parameters to a velocity threshold of 22 degrees/sec, acceleration threshold of 4000 degrees/sec^2^, and travel distance of <1 visual degree. Using these criteria, on average, each participant had 522 trials with at least one microsaccade. To test whether the relationship between theta phase and behavioral performance was attributable to these small, fixational eye movements (i.e., microsaccades), we re-calculated phase-behavior relationships (see previous section) after excluding these trials (Fig. S2A). After excluding trials with microsaccades, there were an average of 376 trials remaining per subject. It is important to note that the measured strength of phase-behavior relationships depends on trial counts (as do other phase-based coherence measures, such as inter-trial phase coherence). Therefore, the scales of results with and without microsaccades (Fig. S2A vs. Fig. S2B) are not comparable (i.e., due to lower trial counts in the analyses without microsaccades). This control analysis revealed that the pattern of phase-behavior results is similar, regardless of whether trials with microsaccades are included or excluded from the analyses.

### Probe-locked event related potentials (ERP) and centro-parietal positivity (CPP)

To assess neurophysiological differences between conditions, we calculated the grand-average voltage locked to the probe onset. For all analyses, we used a baseline from 0.1 to 0s before the probe onset. While both the external and internal probes elicited notable potentials, we focused our analyses on the external probe due to the behavioral effects that we observed in association with these conditions (Fig. 3, Fig. 4). In response to external probes, we identified two primary components: the N1 and the CPP. The N1 is associated with the sensory response to a visual stimulus and is enhanced (more negative) with visual attention to the stimulus^66^. In our case, we observed a slightly late external probe N1 peak (~230 ms, see Fig. 7), presumably due to the low-contrast nature of the external probe. Not shown is the response to the full-contrast internal probe stimulus, which exhibited an N1 peak around 160 ms. In Figure 5, we show the difference in external probe N1 when there is a “bad” pre-probe phase versus a “good” pre-probe phase (see “good versus bad theta bin analyses” section). The other component that we observed in response to the external probe, the centro-parietal positivity (CPP), is associated with the accumulation of evidence towards a decision^67, 68^. In our task, participants must decide whether a probe stimulus corresponds to “horizontal” or “vertical.” We observed that the CPP in our task rises starting at 200 ms after the external probe onset and continues to rise for 200-500 ms afterwards (Fig. 8A, Fig. S4). We show in Figure S4 that CPP latency and magnitude strongly correlate with variability in response times. This supports the previous interpretation of this component as a marker of ongoing decision making^67, 68^. In Figure 8, we show the difference in “match” versus “nonmatch” CPP progression when there is a “bad” pre-probe phase versus a “good” pre-probe phase (see “good” versus “bad” theta bin analyses section).

**Figure 7.**
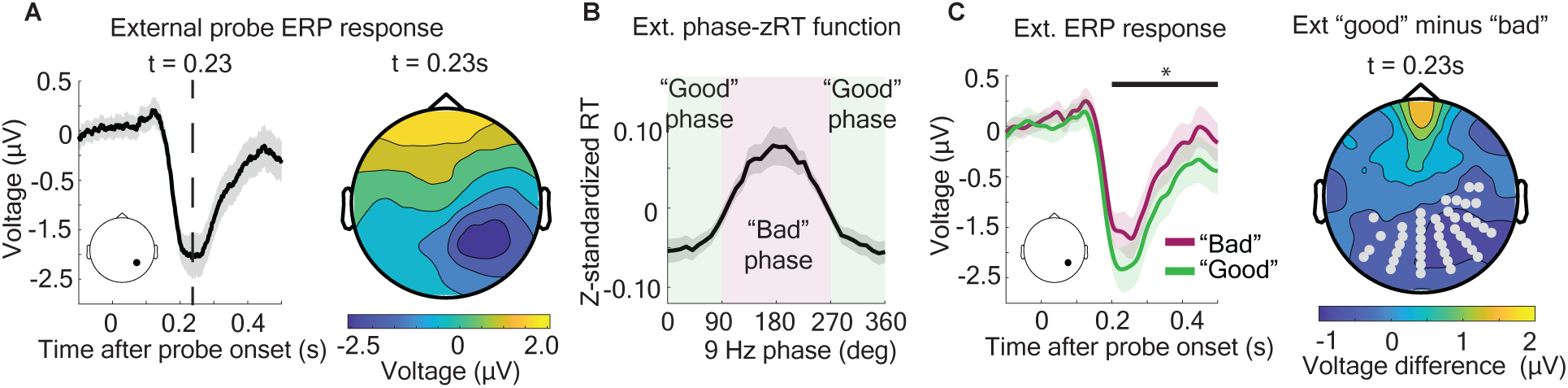
Pre-probe phase affects the probe-locked ERP (on external-probe trials). (A) ERP time-locked to onset of the external probe. *Left:* single-electrode ERP. Inset shows electrode position. *Right:* Topography of a probe-locked ERP at the time of peak negativity (t = 0.23s after the probe). (B) Depiction of the ‘good’ and ‘bad’ pre-probe phases from the phase-RT function (at 4 Hz). (C) *Left:* Single-electrode ERP, depending on whether the pre-probe phase was ‘bad’ (i.e., associated with slower RTs) or ‘good’ (i.e., associated with faster RTs). Inset shows electrode location. *Right*: The difference topography (‘good’ minus ‘bad’ pre-probe phase) at the time of peak visual response (t = 0.23s after probe). All topographical data was flipped to control for probe side. That is, when the external probe was on the right side of fixation, the topography was flipped, effectively making all trials ‘left-probe’ trials. Light gray dots (C, *right*) indicate electrodes that formed a p<0.05 significant spatiotemporal (i.e., across the head and across timepoints) cluster of the voltage difference between the ‘good’ versus ‘bad’ pre-probe phase conditions. Solid line (C, *left*) indicates timepoints for which the inset electrode (included in the cluster in C, *right*) was significant.

**Figure 8.**
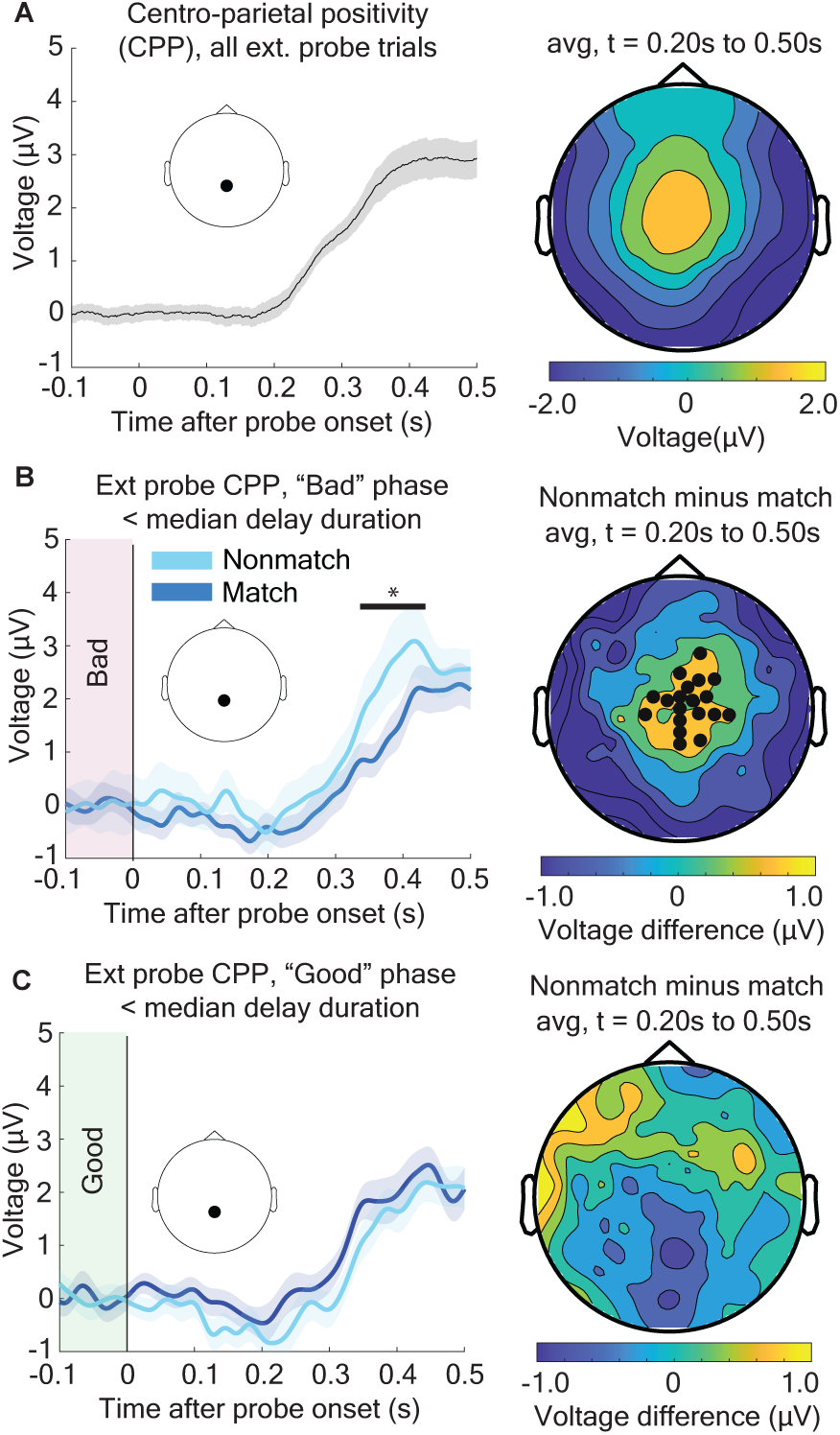
A decision-related ERP component of the memory-interference effect is differentially affected by pre-probe phase. (A) Depiction of the central positivity in response to the low-contrast external probe (consistent with the CPP component, see Figure S5). *Left*, the voltage at a central electrode for all external-probe trials, grand-averaged across participants. Inset shows electrode location. *Right*, topography of grand-averaged voltage from 0.2–0.5s for all external-probe trials. (B) The CPP component for ‘match’ and ‘nonmatch’ trials, specifically for trials when the 4-Hz phase at the frontal electrode was between 90 and 270 degrees (i.e., the ‘bad phase’). (C) Same as (B), but for trials when the 4-Hz phase at the frontal electrode was between 270 and 90 degrees (i.e., the ‘good phase’). Solid bar depicts values belonging to a p<0.05 two-tailed, significant cluster.

For display purposes, we grand-averaged the ERPs across participants and applied a 30-Hz lowpass filter. To test whether ERPs significantly differed between conditions, we used a cluster-based approach to correct for multiple comparisons, across electrodes and timepoints^60^ (see “cluster-based statistics in phase-behavior, windowed behavior, and ERPs” section). All ERP topographies shown (i.e., in Fig. 7, Fig. 8, Fig. S4) have “flipped” electrodes to control for probe side. Specifically, for “probe-side right” trials, the electrodes on the left and right sides of the head were mirrored across the vertical midline, effectively making all trials “probe-side left” trials for these topographies.

“Good” vs. “bad” theta bin analyses

To measure the potential effect of rhythmic dynamics on the interaction between external and internal sampling, we performed some electrophysiological analyses separately for “good” and “bad” theta bins (i.e., theta-phase bins associated with either better or worse behavioral performance). To define these bins, we used the phase-RT function for external sampling at the center of the frontal cluster, at 4 Hz (see Fig. 5), and separated the peak and trough of performance (Fig. 7B). Good-phase trials were defined as trials with pre-probe phases less than 90° and greater than 270°, while bad-phase trials were defined as trials with pre-probe phases greater than 90° and less than 270°.

### Cluster-based statistics in phase-behavior, windowed behavior, and ERPs

We statistically accounted for multiple comparisons and the autocorrelation of behavioral and EEG time-series data over spatial, temporal, and spectral dimensions by applying a nonparametric “cluster-correction” method, which has been widely used in electrophysiological research^13^. Procedurally, this method involves four main steps. First, a distribution of 1500 “null observations” was generated by repeatedly randomizing the data and recalculating the results (see below paragraph for details on how this was specifically done for phase-behavior and windowed behavioral analyses). Second, a distribution of *“*null cluster magnitudes” was created from this distribution of “null observations”. To calculate the magnitude of one null cluster, we first calculated the p-value of each null observation, relative to the rest of the null observations (i.e., the null distribution). Then, the p-values that crossed the significance threshold (p<0.05) were put into “clusters.” The clusters were defined by consecutively adjacent significant values. Adjacency is defined in the spatial domain as electrodes that are directly next to each other on the scalp, in the frequency domain as frequencies that are directly previous or next in sequence (e.g., 4 Hz is adjacent to 5 Hz), and in the temporal domain as timepoints that are directly before or after (e.g., for a 100 Hz sampling rate time series, 0.01s is adjacent to 0.02s, which is adjacent to 0.03s, etc.). Therefore, a cluster represents a grouping of significant values that are unbroken in adjacency across N dimensions. We calculated the magnitude of a cluster by summing 1/p-value of each point in the cluster. This means that a high-value cluster will contain values that have a strong effect (e.g., higher phase-behavior strength or a greater difference in accuracy or voltage) and cover many points in the N-dimensional range of points. After calculating the null clusters for one null observation, the magnitude of the largest null cluster was recorded. For 1500 permutations from the first step, this means 1500 null cluster magnitudes assuming the null hypothesis. Third, the real cluster magnitudes were calculated. This follows the same procedure as step two above, except the real observations are clustered against the null distribution of observations. Once again, this results in a grouping of p<0.05 significant values that have unbroken adjacence. Fourth, the real cluster magnitudes are compared against the null distribution of cluster magnitudes. This resulted in each observed cluster receiving its own p-value compared to the distribution of null-clusters. For cluster-corrected analyses, only the observed values within clusters of p<0.05 magnitude were treated as significant. In summary, a significant value within a significant cluster indicates that not only was this value p<0.05 significant compared to random permutations, but that the magnitude and number of consecutively adjacent values was also significant compared to the null hypothesis.

In the first step described above, a null distribution of observations is needed to create the null clusters which will be compared against the observed clusters. The null hypothesis for phase-behavior analyses is distinct from behavioral and ERP analyses in which the condition labels are simply shuffled. For phase-RT relationships (Fig. 5), the null hypothesis is that there is no relationship between phase and response times. The null distribution is therefore generated by randomizing trial-level phase measurements relative to trial-level RTs (i.e., by breaking the potential relationship between phase and RTs).

## RESULTS

Here, we tested whether and how the brain temporally coordinates shared neural resources associated with selective sampling of the external environment and selective sampling of internal memory stores. We considered two models of temporal coordination (Fig. 1). The first model predicts that external and internally stored information (i.e., an external stimulus in space and/or a memorized stimulus in working memory) are selectively sampled (or enhanced) at different *phases* of a theta rhythm (Fig. 1A). This model is consistent with previous evidence that neural activity associated with different to-be-remembered items can be associated with different phases within an oscillatory cycle, thereby helping to prevent representational conflicts^51–53, 55^. The second model predicts that external information and internally stored information are selectively sampled at different *cycles* of attention-related oscillatory activity (Fig. 1B). This model is consistent with the ‘Rhythmic Theory of Attention’, which proposes that selective processing is characterized by alternating attentional states associated with either sampling at the present focus of attention or an increased likelihood of shifting attentional resources^40^. During the ‘shifting state’, attentional resources might shift from enhancing behaviorally important, external information to enhancing behaviorally important, internally stored information (or vice versa). The second model (Fig. 1B) is also consistent with the conceptualization of internal sampling as internal selective attention^5–11^.

To provide evidence for or against these models, we designed an experimental task that included conditions of (i) only external sampling, (ii) only internal sampling, or (iii) both external and internal sampling (Fig. 2A). Participants maintained fixation throughout the duration of each trial and indicated, following a variable delay (0.3-2.5 seconds), whether a pre-cued visual target (i.e., ‘external-probed’ trials) or a retro-cued memory item (i.e., ‘internal-probed’ trials) was either a horizontal grating or a vertical grating. On non-dual-task (‘alone’) trials, cue/memory stimuli were only presented for one modality (only pre-cues or only memory items). On dual-task (‘both’) trials, cue/memory stimuli were presented for both modalities and either a visual target or a retro-cue could occur, with equal probability (i.e., participants could not anticipate whether a dual-task trial would conclude with an external probe or an internal probe). Dual-task trials included three subtypes: (i) external and internal sampling at the same one location (i.e., ‘both, 1L same’), (ii) external and internal sampling at different locations (i.e., ‘both, 1L diff’), and (iii) external and internal sampling at the same two locations (i.e., ‘both, 2L same’). To promote the deployment of selective attention in anticipation of visual targets, we added noise to the visual gratings (Fig. 2A), pinning each subject’s accuracy on external-only trials at approximately 80 percent. In comparison, accuracy on external-probed, dual-task trials was allowed to vary from 80 percent (i.e., to measure changes in behavioral performance associated with dual-task trials). Memory items—presented prior to the variable delay—were noiseless visual gratings.

We first measured whether dual-task trials were associated with differences in accuracy and response times (RTs) relative to external-only trials and internal-only trials. Previous research has shown that interactions between ongoing perception and working memory decrease over time (i.e., at longer delays), perhaps reflecting a transition from initial working memory representations in sensory regions to more abstract representations in higher-order regions^16^. We therefore took delay length into account for the behavioral analyses. When splitting external-probed trials by median delay (i.e., trials with a delay less than 1.4s and trials with a delay longer than 1.4s), a repeated measures ANOVA revealed accuracy effects of cue/memory condition (levels: ‘alone’, ‘1L same’, ‘1L diff’, ‘2L same’; F = 4.22 and p = 0.012), delay duration (levels: less than median, longer than median; F = 30.946 and p < 0.0001), and an interaction between cue/memory condition and delay duration (F = 6.63, p = 0.0014) (Fig. 3A, B). Follow-up two-tailed t-tests showed that external-only (‘alone’) trials had significantly higher accuracy than ‘1L same’ (p = 0.0089) and ‘2L same’ (p = 0.0044) trials at short delays (Fig. 3A). RTs to the external probe, like accuracy, showed effects of cue/memory condition (F = 14.463 and p < 0.0001), delay duration (F = 53.47 and p < 10^-6^), and an interaction between condition and delay duration (F = 5.20 and p = 0.0037) (Fig. 3D, E). At delays shorter than the median (1.4s), RTs were significantly faster for external-only (‘alone’) trials than ‘1L same’ (p = 0.042) and ‘2L same’ (p = 0.012) trials. Unlike with external-probed trials, a repeated measures ANOVA showed no significant effects for internal-probe accuracy (Fig. S1A, B) or internal-probe RTs (Fig. S1D, E).

To better visualize the time-course of interactions between external and internal sampling (relative to the length of the delay), we plotted time-windowed behavioral performance (Fig. 3B, 3D, S1C, S1F), using overlapping 600-ms bins. That is, we plotted behavioral performance as a function of delay length. There were significant clusters for external-probe time-windowed accuracy (p = 0.00055) and RTs (p = 0.0094) (Fig. 3C, F). Interference effects (i.e., the effects of working memory on the processing of an external probe) persisted for between 1.2–1.5s after the offset of the flash stimulus (i.e., 1.55–1.85s after the offset of the pre-cue and memory items). Consistent with previous research, these behavioral analyses suggest that external and internal sampling share neural resources, with interactions between ongoing perception and working memory only occurring during shorter delays^16^.

We next took a closer look at interactions between external and internal sampling. Previous research has demonstrated that interactions between external and internal sampling can depend on the degree of similarity between environmental information and internally stored information^16, 17^. Here, we binned external-probed, dual-task (‘both’) trials based on whether the to-be-detected information (i.e., the low-contrast grating) was either a ‘match’ or a ‘non-match’ for the to-be-remembered information (i.e., the memory item). For the ‘both, 1L same’ condition and the ‘both, 2L same’ conditions (see Fig. 4A), a match was defined by both the orientation (i.e., whether the gratings were horizontal or vertical) and the spatial location of the gratings (i.e., whether the matching information occurred at the same location in space). For the ‘both, 1L diff’ condition, however, a match was only defined by the orientation of the gratings (i.e., the external and internal information for this condition always occurred at different spatial locations). As with the behavioral plots in Figure 3, we split the trials based on the median delay into short-delay trials (delays < 1.4s) and long-delay trials (delays >= 1.4s) (Fig. 4B). A repeated measures ANOVA showed main effects of cue/memory condition (levels: ‘1L same’, ‘1L diff’, ‘2L same’; F = 3.44 and p = 0.040) and delay duration (levels: ‘less than median’, ‘longer than median’; F = 39.04 and p < 10^-5^) and a significant two-way interaction of cue/memory condition with delay duration (F = 4.94, p < 0.036) and a significant three-way interaction of cue/memory condition with delay duration and match status (F = 3.71, p = 0.032). Follow-up two-tailed t-tests revealed significantly higher accuracy for ‘nonmatch’ relative to ‘match’ trials for the ‘both, 1L same’ (p = 0.0410) and ‘both, 2L same’ (p = 0.0365) conditions but not the ‘both, 1L diff’ condition (p = 0.8013). These results indicate that ‘match effects’ only occurred during shorter delays and were dependent on both matching orientation and matching spatial location.

To better visualize the time-course of match effects, we also calculated accuracy for ‘match’ and ‘non-match’ trials over time-windowed delay bins (i.e., overlapping 600-ms bins). Here, we restricted our analysis to the conditions previously associated with significant match effects (i.e., the ‘both, 1L same’ and ‘both, 2L same’ conditions). Figure 4C indicates that match effects persisted for approximately 1.4s after the delay onset, consistent with our previous analysis of interactions between external and internal sampling (Fig. 3). These match effects (or interference effects) partly explain the decrement in behavioral performance for external probes when comparing ‘external-alone’ trials with ‘external-both’ trials (Fig. 3).

After establishing behavioral interactions between external and internal sampling within the context of the present experimental task, we tested whether and how external and internal sampling share a theta-rhythmic sampling process (i.e., our primary research question)^40^. Here, we calculated phase-behavior relationships, which are a quantification of how pre-probe phase predicts behavioral performance (Fig. 5). We used wavelets to measure frequency-specific phase (from the EEG signal) during the variable delay, just prior to either a visual target (i.e., during external-probe trials) or a retro-cue (i.e., during internal-probe trials)^44, 51, 63, 64^. Because accuracy on internal-probe trials was near ceiling, we focused on the relationship between frequency-specific phase (from 3–55 Hz) and RTs (but see later analyses showing that accuracy for external-probe trials similarly fluctuates as a function of theta phase, Fig. S3. After obtaining pre-probe phase measurements for each trial at each electrode, we calculated median RTs in overlapping phase bins (i.e., we measured RTs as a function of phase, see Fig. 5D). We then fit the phase-RT functions with a one-cycle sine wave and used the amplitude of that sine wave to measure the strength of the phase-RT relationships (after averaging the phase-RT functions across participants)^44, 51, 63, 64^. This approach is based on the assumption that there will be a phase associated with relatively better behavioral performance, and 180 degrees from that phase, a phase associated with relatively worse behavioral performance^44, 63^. Figure 5A shows the strength of the phase-RT relationships for every electrode (1–128) and frequency (3–55 Hz) combination, separately for external- and internal-probed trials (after combining the ‘only’ and ‘both’ conditions). Here, we zeroed statistically insignificant results, based on a non-parametric permutation approach and cluster-based statistics (i.e., to correct for multiple comparisons)^60^. Figure S2B shows the non-zeroed results. The frequencies and electrodes associated with significant phase-RT relationships (p < 0.05) were strikingly similar across external- and internal-probed trials (Fig. 5A), with significant phase-RT relationships primarily occurring in the theta- and alpha-frequency bands, peaking at 4–5 Hz and 9–11 Hz (Fig. 5B). Figure 5C shows the clusters of electrodes with significant phase-RT relationships and the associated topographies at 4 Hz and 9 Hz. Because the likelihood of microsaccades has also been shown to fluctuate as a function of theta phase^65^, we re-calculated our results after excluding trials with microsaccades occurring during the delay. Supplemental Figure 2A, consistent with previous research^44, 46^, shows that the pattern of phase-RT relationships is similar with and without microsaccades, indicating that phase-RT relationships are not attributable to microsaccades.

In addition to examining whether significant phase-RT relationships for external and internal sampling occurred at similar electrodes and frequencies, we measured whether faster and slower RTs for external and internal sampling were associated with similar phases (Fig. 5D)^51^. Here, we used phase-RT functions from a single electrode, taken from the frontal cluster (i.e., the electrode with the strongest phase-RT relationship). As shown in Figure 5D, the phases associated with faster and slower RTs, at 4Hz, were consistent for external- and internal-probed trials. We used a circular Watson-Williams test^51, 69^ to statistically test whether the specific phase associated with faster RTs differed between external- and internal-probed trials. This test demonstrated that the distributions of fast-RT phases (across participants) for external- and internal-probed trials were not statistically different between these conditions (p = 0.2435). Figure 5E shows the circular histograms associated with fast-RT phases for external- and internal-probed trials (i.e., the distributions of subject-specific, fast-RT phases).

Figure 6 shows condition-specific phase-RT relationships for external- and internal-probed trials, i.e., separately for ‘only’ trials and each subtype of the dual-task conditions (i.e., ‘both, 1L same,’ ‘both, 1L diff,’ and ‘both, 2L same’). Here, we show these relationships at 4 Hz (note: the results at 4 Hz and 9 Hz are highly consistent, as shown in Fig. 5). While the significant electrode clusters vary across these conditions, the topographies (Fig. 6A) and the specific phases (Fig. 6B,C) associated with faster and slower RTs are similar. That is, the results are largely similar for external- and internal-probed trials, regardless of whether trials required only external sampling, only internal sampling, or both external and internal sampling. We again used a circular Watson-Williams test^51, 69^ to measure whether the specific phases associated with faster RTs differed between external- and internal-probed trials (Fig. 6C). This test demonstrated that the distributions of fast-RT phases (across participants) for external- and internal-probed trials were not statistically different between these trial types for ‘only’ trials (p = 0.5121), ‘both, 1L same’ trials (p = 0.9934), ‘both, 1L diff’ trials (p = 0.0798), or ‘both, 2L same’ trials (p = 0.1650). Rather than being coordinated at different phases of oscillatory activity, the present results are consistent with a rhythmic process that selectively samples either external or internal information at a shared peak behavioral phase (i.e., the “cycles” model in Fig. 1B).

In addition to phase-RT relationships, we calculated phase-accuracy relationships (Fig. S3. Due to ceiling effects for internal-probe accuracy (>90%), we only used the external-probe condition. We found that the strength of phase-accuracy relationships created a highly similar frequency/electrode cluster (Fig. S3A) to phase-RT relationships (Fig. 5). Likewise, the topography of the phase-accuracy relationships (Fig. S3B) was comparable to the topography of the phase-RT relationships (Fig. 5C). Furthermore, the phase associated with the highest accuracy (Fig. S3C, D) was the same as the phase associated with the fastest RTs (Fig. 5D, E).

Across all conditions, there was a pre-probe phase associated with better behavioral performance (i.e., faster, more accurate behavioral performance) and a pre-probe phase associated with worse behavioral performance (i.e., slower, less accurate behavioral performance) (Fig. 7B). These ‘good’ and ‘bad’ phases are approximately 180 degrees apart, and therefore the 360-degree span of phases can be used to split trials into two bins: ‘good-phase’ trials and ‘bad-phase’ trials (Fig. 7B). To investigate how these pre-probe phases are linked to differences in behavioral performance, we calculated probe-locked ERPs (Fig. 7A), separately for these two types of trials (i.e., trials binned by ‘good’ or ‘bad’ pre-probe phase, Fig. 7C). We found that there was a significant spatiotemporal (i.e., across electrodes and time) cluster (p < 0.001) of posterior electrodes where the probe-evoked initial negativity (i.e., the N1 component^66^), contralateral to the probe, was significantly stronger on ‘good-phase’ trials relative to ‘bad-phase’ trials (Fig. 7C). These results indicate that pre-probe phase was associated with the subsequent strength of the probe-evoked visual response (on external-probe trials), with stronger probe-evoked visual responses on trials associated with better behavioral performance.

Finally, we tested whether the ‘good’ and ‘bad’ pre-probe phases had differential effects on neurophysiological interactions between external and internal sampling, specifically by further binning same-location ‘match’ and ‘non-match’ trials (see Fig. 4) based on pre-probe phase (i.e., based on whether there was a ‘good’ or a ‘bad’ phase prior to the occurrence of the external probe). Again, a ‘match’ occurred when the to-be-detected information (e.g., a horizontal grating in noise) on an external-probe trial matched the to-be-remembered information. We specifically measured whether probe-evoked ERPs were different across memory interference conditions (i.e., ‘match’ vs. ‘nonmatch’) associated with either a ‘good’ pre-probe phase or a ‘bad’ pre-probe phase (Figure 8B, C). The results revealed a significant cluster (p = 0.006) of spatiotemporal (over electrode and time) differences between match and nonmatch trials when the pre-probe phase was ‘bad’ (Fig. 8B), but not when the pre-probe phase was ‘good’ (Fig. 8C). These differences were consistent with a modulation of the centro-parietal positivity (CPP): an ERP component that is associated with the accumulation of perceptual and decision-related evidence (Fig. 8A)^67, 68^. Supplemental Figure 4 demonstrates that both the magnitude and the latency of the CPP were dependent on RTs, presumably reflecting the accumulation of evidence towards a decision (i.e., either vertical or horizontal)^67, 68^. The results shown in Figure 8 indicate that interactions between external and internal information (e.g., interference associated with matching external and internal information) can fluctuate with the pre-probe phase of rhythmic sampling processes.

## DISCUSSION

Everyday tasks require the selective sampling (i.e., the attention-related sampling) of behaviorally important information, both environmental information and internally stored information (e.g., information being maintained in working memory). Recent evidence suggests that aspects of this selective sampling, whether turned outward^40–50^ or inward^54, 56^, fluctuate at a theta frequency (3–8 Hz). In the context of external sampling (i.e., from the environment), attention-related behavioral and neural effects wax and wane as a function of theta phase^40–50^. We have proposed that these temporal dynamics reflect alternating attentional states associated with either sampling (i.e., sensory functions of the large-scale network that directs selective attention) or a greater likelihood of shifting (i.e., motor functions of the large-scale network that directs selective attention)^40, 41, 44^. These alternating attentional states^44, 64^ might help to resolve *functional* conflicts attributable to shared neural resources. In the context of internal sampling (i.e., from internal memory stores), theta-rhythmic neural activity might also, for example, help to resolve *representational* conflicts attributable to shared neural resources. Specifically, previous work has shown that the strength of neural representations associated with to-be-remembered items waxes and wanes as a function of theta phase^51–56^ (or theta-dependent, higher frequency phase^51, 52^), with different to-be-remembered items being associated with different phases (i.e., there is a phase-specific coding of different items). In other words, whether a task involves the selective sampling of environmental information or the selective sampling of internally stored information, the selection and/or enhancement of behaviorally important information fluctuates at a theta frequency.

Whereas previous work has separately investigated theta-rhythmic, attention-related sampling of environmental and internally stored information, we tested whether there is a domain-general process for the selective sampling of behaviorally important information. That is, we investigated whether theta-rhythmic sampling of environmental and internally stored information share neural resources. Here, we considered two potential proposals for how external and internal sampling might be coordinated relative to oscillatory phase, given a shared neural mechanism for theta-rhythmic sampling: (i) the selective enhancement of external information and the selective enhancement of internal information might alternate within a theta cycle (Fig. 1A) or (ii) the selective enhancement of external information and the selective enhancement of internal information might alternate across different theta cycles (Fig. 1B). The second proposal would be consistent with ‘The Rhythmic Theory of Attention,’ with theta-dependent increases in the likelihood of attentional shifts (i.e., the ‘shifting state’) being associated with either within-domain shifts (e.g., across different locations in the visual field) or between-domain shifts (e.g., from external information to internal information).

To test these hypotheses, we compared evidence of theta-rhythmic sampling across trials that required (i) only external sampling, (ii) only internal sampling, or (iii) both external and internal sampling (i.e., dual-task trials). Given a shared neural mechanism for theta-rhythmic sampling, dual-task trials would create a competition for limited processing resources. Prior to testing for theta-rhythmic sampling, we measured behavioral interactions between external and internal sampling on dual-task trials relative to external- and internal-only trials (Figs. 3 and 4). These behavioral interactions provided indirect evidence of shared neural resources, consistent with previous research^29, 32–39^. We next demonstrated that theta phase prior to the onset of either a visual target (i.e., during external-probe trials) or a retro-cue (i.e., during internal-probe trials) was linked to similar fluctuations in behavioral performance (Figs. 5 and 6), regardless of whether there was a potential conflict between external and internal sampling (i.e., during dual-task trials) or not (i.e., during external-only trials or internal-only trials). Both the topographies of behaviorally relevant theta-band activity (i.e., phase-RT relationships) and the specific theta phases associated with either better or worse behavioral performance were similar across trials that probed external sampling and trials that probed internal sampling (Figs. 5 and 6). The present findings are therefore consistent with a shared theta-rhythmic process for enhancing either environmental information or internally stored information. Given the spatial limitations of the EEG signal, attributable to volume conduction, future research—using methods with greater spatial resolution (e.g., neurophysiology in non-human primates)^7, 36, 37^—will need to address whether this shared sampling process (i.e., domain-general sampling process) (i) can concurrently sample external and internal information, perhaps by interacting with domain-specific neural populations^36, 37^, or (ii) alternates between periods of sampling external information and periods of sampling internal information (Fig. 1B). To clarify the second alternative, it may be that the shared process for theta-rhythmic sampling can either be turned inward or outward at any given moment in time (but not concurrently inward and outward).

While definitively establishing temporal isolation of attention-related external sampling from attention-related internal sampling will require further research, consistency in the theta phases associated with either better or worse behavioral performance (Figs. 5D and 6B) indicates that external and internal sampling are not being temporally coordinated within a single theta cycle (Fig. 1A). That is, there was no apparent difference between the phases associated with better behavioral performance for external versus internal sampling. If theta-rhythmic enhancement of external information and theta-rhythmic enhancement of internal information are occurring during different time periods (rather than concurrently), switches between these processes must occur between theta cycles (Fig. 1B).

Future research will also need to test whether there is a specific theta phase associated with a higher likelihood of shifts (or switches) between periods of bias toward either external sampling or internal sampling. Although between-domain shifts (e.g., from external to internal) have yet to be examined in the context of theta-rhythmic sampling, previous work has provided evidence that within-domain shifts (e.g., attentional shifts between different locations in the environment) occur at a consistent theta phase^40, 48, 58, 70^. Evidence for or against a common mechanism for shifting within or between domains is mixed^15^. Most research has separately investigated shifting within the contexts of either external sampling or internal sampling (i.e., focusing exclusively on within-domain shifts). Recent research, however, has shown (i) that between-domain shifts lead to greater behavioral (or switch) costs than within-domain shifts^32, 71^, and (ii) that a multivariate classifier can discriminate within-from between-domain shifts^32^. These findings suggest at least partly distinct underlying neural processes. The same research, however, also demonstrated that changes in alpha-power lateralization attributable to within- and between-domain shifts have equivalent time courses^32^. Alpha-power lateralization is associated with spatial orienting^72^, and equivalent time courses for changes in alpha-power lateralization (i.e., when the to-be-sampled information shifts from one visual hemifield to the other) provide evidence that within- and between-domain shifts have equivalent time courses. Based on evidence from the present experiment (Figs. 5 and 6) of a shared, theta-rhythmic process for selective sampling of either external or internal information, we predict that the likelihood of between-domain shifts—like within-domain shifts^40, 48, 58, 70^—fluctuates as a function of theta phase.

While we have focused on the relationship between theta phase and behavioral performance (peaking at 4–5 Hz), phase-RT relationships in the present experiment spanned lower frequencies, with a second peak in the alpha band (at 9–11 Hz). The topographies of phase-RT relationships in the theta and alpha bands were highly consistent (Fig. 5C), suggesting that these effects reflect the same neural generators. The presence of phase-behavior relationships across multiple frequencies is consistent with previous findings^44, 51, 63, 64^. We have previously linked different behaviorally relevant frequencies, for example, to different functionally defined cell types (via spike-LFP phase coupling)^44^. Future research (e.g., using intracortical recordings in non-human primates) will need to better parse the functional overlap and/or segregation between theta- and alpha-band activity (i.e., the specific functions associated with behaviorally relevant theta- and alpha-band activity).

Finally, we tested whether theta phase modulates (i) target-evoked visual responses (on external-probe trials) and (ii) neurophysiological evidence of interactions between external and internal sampling. The theta phase associated with better behavioral performance (i.e., the ‘good’ phase) was also associated with stronger target-evoked visual responses, onsetting during the N1 component of the ERP (Fig. 7). For dual-task trials (i.e., ‘both’ trials), the theta phase associated with worse behavioral performance was also associated with a slower accumulation of perceptual evidence for decision making (as measured via the CPP component), specifically when to-be-detected information matched to-be-remembered information (Fig. 8). A shared, theta-rhythmic sampling process can therefore modulate interactions between environmental information and internally stored information during dual-task trials, when these sources of information compete for limited processing resources. Theta-rhythmic sampling influences both selective sensory processing and subsequent cognitive processing (i.e., evidence accumulation and decision making), leading to phasic fluctuations in behavioral performance (e.g., RTs).

The present findings demonstrate that theta-rhythmic sampling reflects a more general process for selecting and boosting behaviorally important information, regardless of whether that information is sampled from the environment or sampled from internal memory stores. That is, there is a domain-general, theta-rhythmic process—with a shared neural basis—for periodically enhancing either external or internal information. While this process is shared between external and internal sampling, there are likely other aspects of external and internal sampling that are domain specific^32, 73, 74^. Future research will need to (i) determine the neural circuits through which this sampling process interacts with neural representations of environmental information and internally stored information, and (ii) determine the extent to which environmental information and internally stored information can be concurrently sampled through attention-related mechanisms.

## ACKNOWLEDGEMENTS

This work was supported by grants from the National Science Foundation (NSF 2120539) and the Searle Scholars Program to I.C.F, and from the National Institutes of Health (NEI T32EY007125) to P.C. We would like to thank Miral Abdalaziz, Amber McFerren, and Jordan Klembczyk for their help with data collection.

---

**Supplementary Figure 1.**
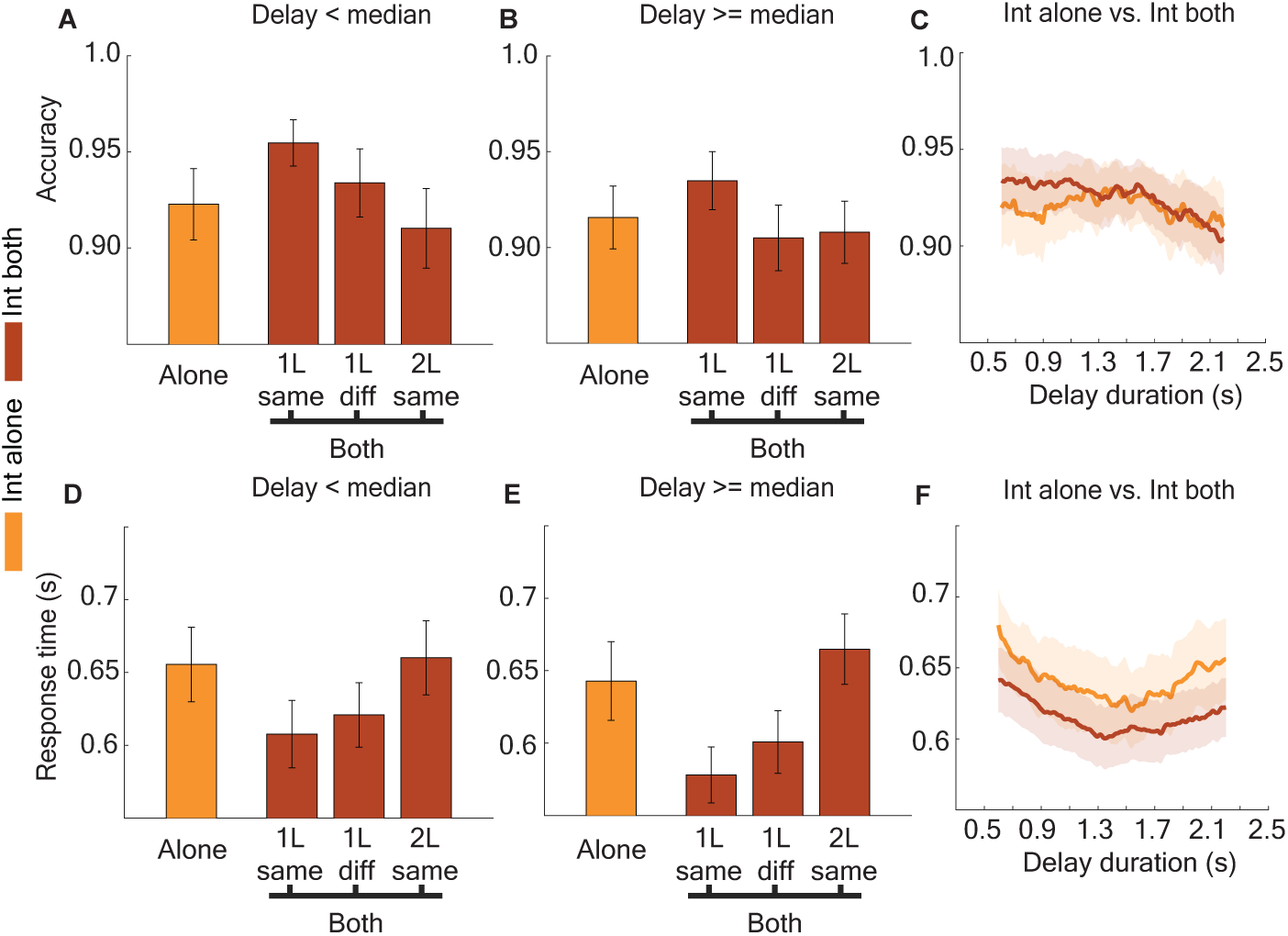
Internal sampling is robust to concurrent perceptual sampling. (A) Accuracy for internal-only and internal-both (“1L same”, “1L diff”, and “2L”) trials in which the delay duration was less than the median (< 1.4s). (B) Same as (A), but for delays longer than the median (>1.4s). (C) Delay-duration windowed accuracy for the external-only and external-both conditions (here, the ‘both’ conditions have been combined). Window length was 0.6s, window step was 0.01s. (D-F) Same as (A-C), but for response times (RTs).

**Supplementary Figure 2.**
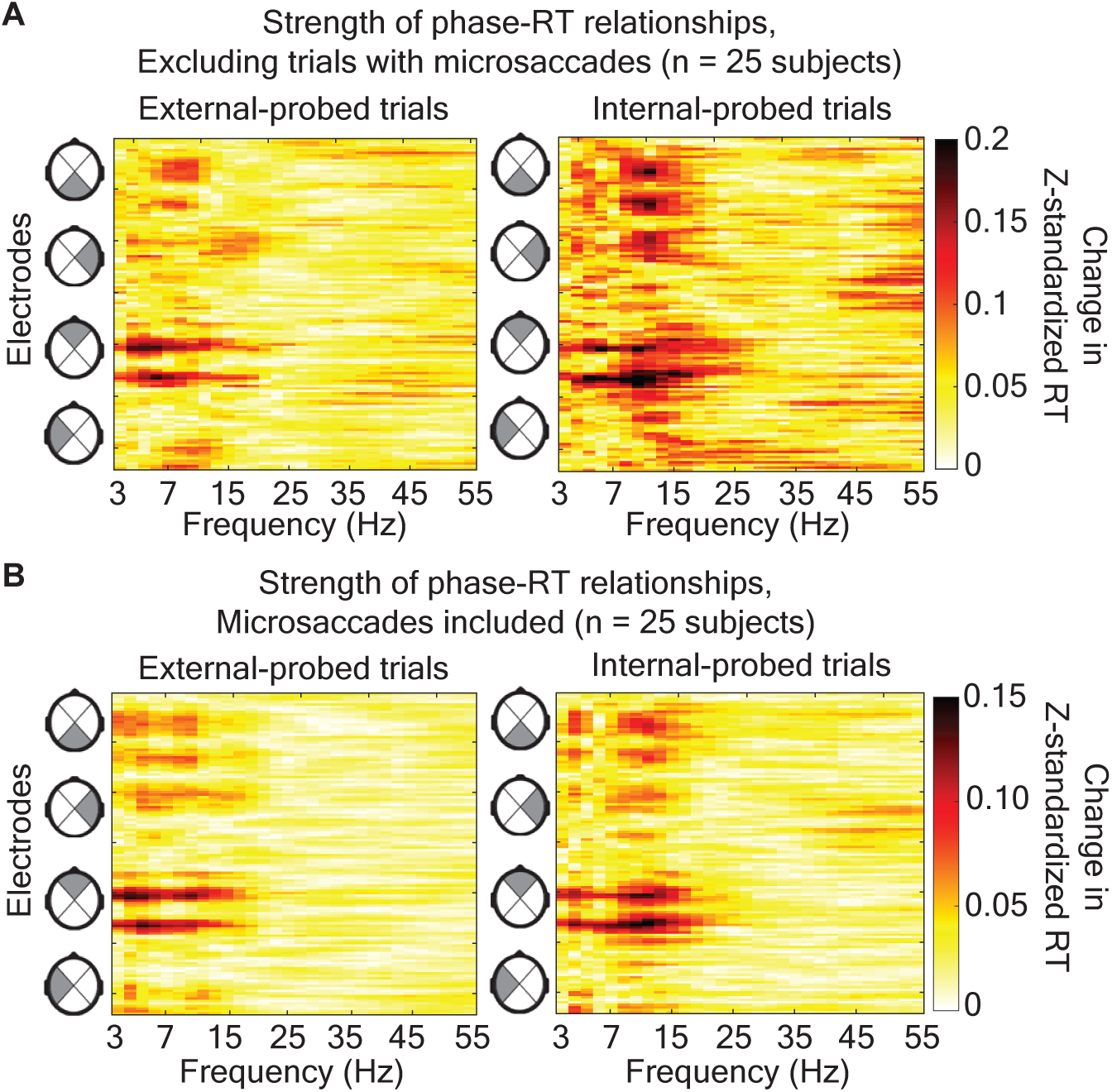
Phase-RT relationships remain consistent after excluding trials with microsaccades. (A) Phase-RT relationships for all electrode (1–128) and frequency (3–55 Hz) combinations, separately for external- and internal-probe trials, after excluding trials with microsaccades. (B) Phase-RT relationships for all electrode and frequency combinations, separately for external- and internal-probe trials, including trials with microsaccades (same data as Fig. 4A), but without zeroing the insignificant values.

**Supplementary Figure 3.**
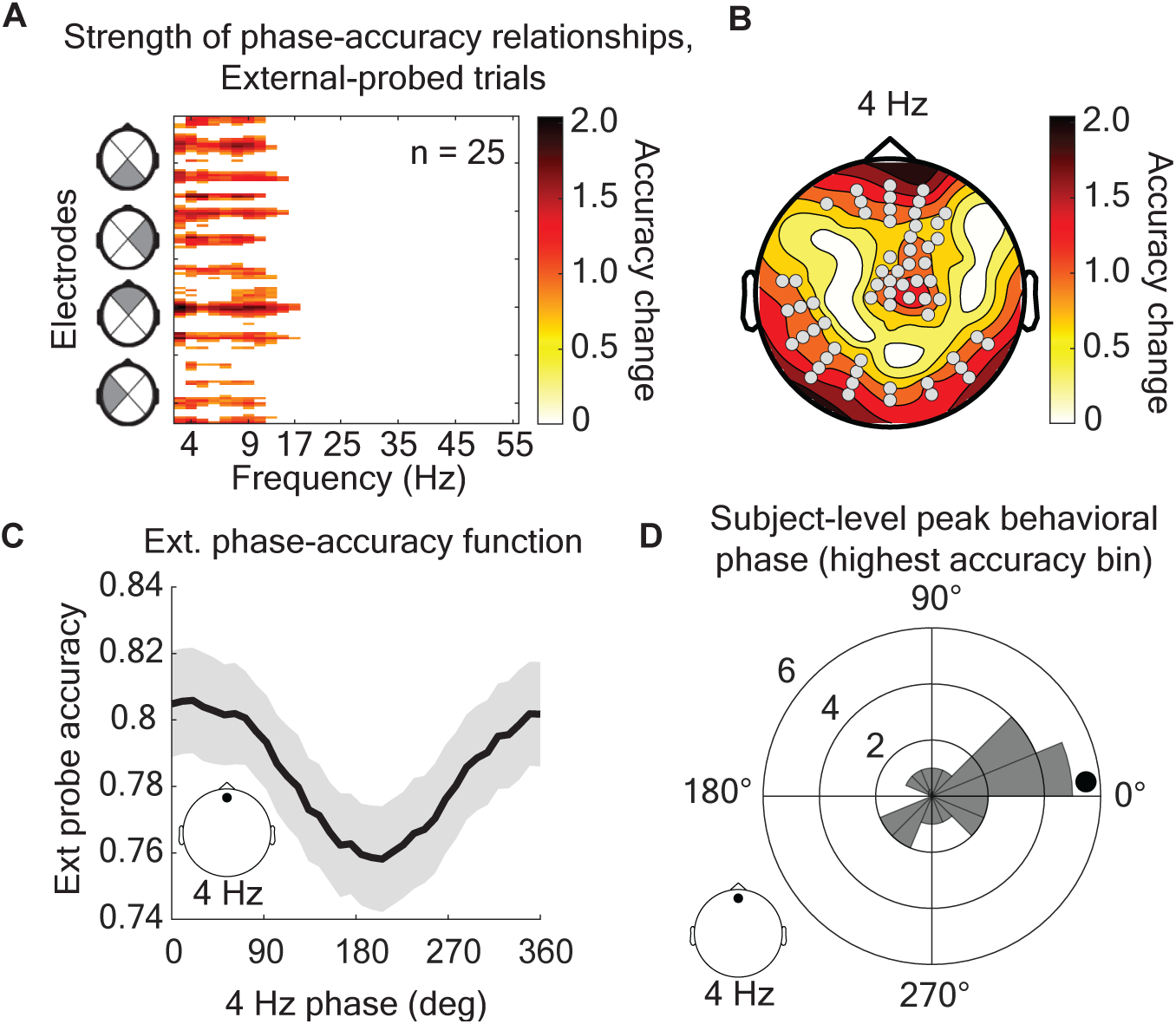
Phase-accuracy relationships show a similar pattern as the phase-RT relationships. (A) Strength of phase-accuracy relationships are shown for a range of frequencies (3 to 55 Hz) and electrodes (128 total), for all external-probe trials. All values that were not part of p<0.05 significant clusters have been zeroed. (B) Topography of phase-accuracy strength at 4 Hz. Dots depict electrodes for the plotted frequency (4 Hz) that were part of a significant cluster in (A). (C) Grand-averaged phase-accuracy function for all external-probe trials. Note that the phase-accuracy function appears flipped relative to the phase-RT function due to accuracy and RTs having an inverse relationship. Inset represents the electrode location (frontal) and frequency (4 Hz). (D) The phase at which external probe accuracy was best (highest) for each participant. The black circle depicts the grand-averaged, peak phase (i.e., represents the peak phase of the data in C). The radial axis depicts number of participants in each peak-phase bin. Inset represents electrode location (frontal) and frequency (4 Hz).

**Supplementary Figure 4.**
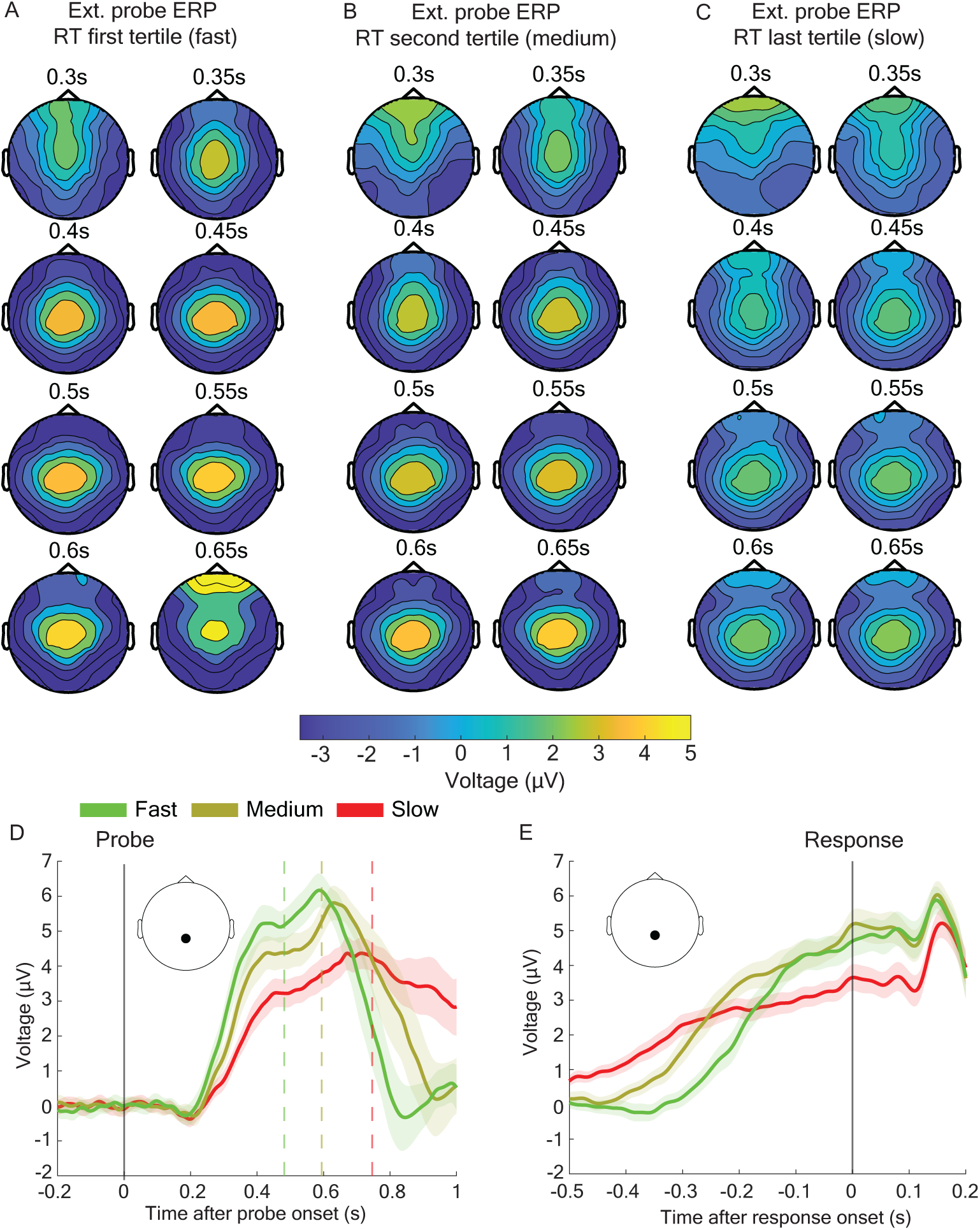
An ERP component categorized as the centro-parietal positivity (CPP) predicts behavioral performance via its latency and magnitude. (A-C) Topographical time-series of the grand-averaged voltage from 0.3 to 0.65s, time-locked to the onset of the external probe. *Left*, fastest tertile of RTs (only correct trials included). *Middle*, middle tertile of RTs. *Right*, slowest tertile of RTs. There is notably a lower magnitude and a longer latency with slower responses to the low-contrast external probe. (D) Probe-locked CPPs for a representative electrode (shown in inset), separately for the three RT tertiles. Colored dashed lines indicate mean RT (relative to probe onset) for the corresponding tertile. Error bars depict SEM of the grand-averaged voltage. (E) Response-locked CPPs for a representative electrode (shown in inset), separately for the three RT tertiles. Error bars depict SEM of the grand-averaged voltage.

## REFERENCES

1. Carrasco M. Visual attention: The past 25 years. Vision research. 2011;51(13):1484–525. doi: 10.1016/j.visres.2011.04.012; PMCID: PMC3390154.

2. Moore T, Zirnsak M. Neural Mechanisms of Selective Visual Attention. Annual review of psychology. 2017;68:47–72. doi: 10.1146/annurev-psych-122414-033400. PubMed PMID: 28051934.

3. D’Esposito M, Postle BR. The cognitive neuroscience of working memory. Annual review of psychology. 2015;66:115–42. Epub 20140919. doi: 10.1146/annurev-psych-010814-015031. PubMed PMID: 25251486; PMCID: PMC4374359.

4. Baddeley A. Working memory. Science. 1992;255(5044):556–9. doi: 10.1126/science.1736359. PubMed PMID: 1736359.

5. Kiyonaga A, Egner T. Working memory as internal attention: toward an integrative account of internal and external selection processes. Psychonomic bulletin & review. 2013;20(2):228–42. doi: 10.3758/s13423-012-0359-y. PubMed PMID: 23233157; PMCID: 3594067.

6. van Ede F, Nobre AC. Turning Attention Inside Out: How Working Memory Serves Behavior. Annual review of psychology. 2023;74:137–65. Epub 20220812. doi: 10.1146/annurev-psych-021422-041757. PubMed PMID: 35961038.

7. van Ede F, Nobre AC. Toward a neurobiology of internal selective attention. Trends in neurosciences. 2021;44(7):513–5. Epub 20210512. doi: 10.1016/j.tins.2021.04.010. PubMed PMID: 33992457.

8. Gazzaley A, Nobre AC. Top-down modulation: bridging selective attention and working memory. Trends in cognitive sciences. 2012;16(2):129–35. doi: 10.1016/j.tics.2011.11.014. PubMed PMID: 22209601; PMCID: 3510782.

9. Myers NE, Stokes MG, Nobre AC. Prioritizing Information during Working Memory: Beyond Sustained Internal Attention. Trends in cognitive sciences. 2017;21(6):449–61. Epub 20170425. doi: 10.1016/j.tics.2017.03.010. PubMed PMID: 28454719; PMCID: PMC7220802.

10. Nobre AC, Coull JT, Maquet P, Frith CD, Vandenberghe R, Mesulam MM. Orienting attention to locations in perceptual versus mental representations. Journal of cognitive neuroscience. 2004;16(3):363–73. doi: 10.1162/089892904322926700. PubMed PMID: 15072672.

11. Theeuwes J, Belopolsky A, Olivers CN. Interactions between working memory, attention and eye movements. Acta Psychol (Amst). 2009;132(2):106–14. Epub 20090223. doi: 10.1016/j.actpsy.2009.01.005. PubMed PMID: 19233340.

12. Chun MM, Golomb JD, Turk-Browne NB. A taxonomy of external and internal attention. Annual review of psychology. 2011;62:73–101. doi: 10.1146/annurev.psych.093008.100427. PubMed PMID: 19575619.

13. Awh E, Jonides J. Overlapping mechanisms of attention and spatial working memory. Trends in cognitive sciences. 2001;5(3):119–26. doi: 10.1016/s1364-6613(00)01593-x. PubMed PMID: 11239812.

14. Oberauer K. Working Memory and Attention - A Conceptual Analysis and Review. J Cogn. 2019;2(1):36. Epub 20190808. doi: 10.5334/joc.58. PubMed PMID: 31517246; PMCID: PMC6688548.

15. Nobre AC, Gresch D. How the brain shifts between external and internal attention. Neuron. 2025. Epub 20250714. doi: 10.1016/j.neuron.2025.06.013. PubMed PMID: 40675155.

16. Teng C, Kaplan SM, Shomstein S, Kravitz DJ. Assessing the interaction between working memory and perception through time. Attention, perception & psychophysics. 2023;85(7):2196–209. Epub 20230922. doi: 10.3758/s13414-023-02785-3. PubMed PMID: 37740152.

17. Teng C, Kravitz DJ. Visual working memory directly alters perception. Nature human behaviour. 2019;3(8):827–36. Epub 20190708. doi: 10.1038/s41562-019-0640-4. PubMed PMID: 31285620.

18. Olivers CN, Meijer F, Theeuwes J. Feature-based memory-driven attentional capture: visual working memory content affects visual attention. Journal of experimental psychology Human perception and performance. 2006;32(5):1243–65. doi: 10.1037/0096-1523.32.5.1243. PubMed PMID: 17002535.

19. Soto D, Heinke D, Humphreys GW, Blanco MJ. Early, involuntary top-down guidance of attention from working memory. Journal of experimental psychology Human perception and performance. 2005;31(2):248–61. doi: 10.1037/0096-1523.31.2.248. PubMed PMID: 15826228.

20. Soto D, Humphreys GW, Heinke D. Working memory can guide pop-out search. Vision research. 2006;46(6-7):1010–8. Epub 20051027. doi: 10.1016/j.visres.2005.09.008. PubMed PMID: 16257030.

21. Rademaker RL, Bloem IM, De Weerd P, Sack AT. The impact of interference on short-term memory for visual orientation. Journal of experimental psychology Human perception and performance. 2015;41(6):1650–65. Epub 20150810. doi: 10.1037/xhp0000110. PubMed PMID: 26371383.

22. Nobre AC, Gitelman DR, Dias EC, Mesulam MM. Covert visual spatial orienting and saccades: overlapping neural systems. NeuroImage. 2000;11(3):210–6. doi: 10.1006/nimg.2000.0539. PubMed PMID: 10694463.

23. Beck VM, Hollingworth A, Luck SJ. Simultaneous control of attention by multiple working memory representations. Psychological science. 2012;23(8):887–98. Epub 20120703. doi: 10.1177/0956797612439068. PubMed PMID: 22760886; PMCID: PMC3419335.

24. Van der Stigchel S, Merten H, Meeter M, Theeuwes J. The effects of a task-irrelevant visual event on spatial working memory. Psychonomic bulletin & review. 2007;14(6):1066–71. doi: 10.3758/bf03193092. PubMed PMID: 18229476.

25. Konstantinou N, Lavie N. Dissociable roles of different types of working memory load in visual detection. Journal of experimental psychology Human perception and performance. 2013;39(4):919–24. Epub 20130527. doi: 10.1037/a0033037. PubMed PMID: 23713796; PMCID: PMC3725889.

26. Konstantinou N, Bahrami B, Rees G, Lavie N. Visual short-term memory load reduces retinotopic cortex response to contrast. Journal of cognitive neuroscience. 2012;24(11):2199–210. Epub 20120820. doi: 10.1162/jocn_a_00279. PubMed PMID: 22905823.

27. Balestrieri E, Ronconi L, Melcher D. Shared resources between visual attention and visual working memory are allocated through rhythmic sampling. The European journal of neuroscience. 2022;55(11-12):3040–53. Epub 20210512. doi: 10.1111/ejn.15264. PubMed PMID: 33942394.

28. Pasternak T, Greenlee MW. Working memory in primate sensory systems. Nature reviews Neuroscience. 2005;6(2):97–107. doi: 10.1038/nrn1603. PubMed PMID: 15654324.

29. Emrich SM, Riggall AC, Larocque JJ, Postle BR. Distributed patterns of activity in sensory cortex reflect the precision of multiple items maintained in visual short-term memory. The Journal of neuroscience : the official journal of the Society for Neuroscience. 2013;33(15):6516–23. doi: 10.1523/JNEUROSCI.5732-12.2013. PubMed PMID: 23575849; PMCID: PMC3664518.

30. Serences JT. Neural mechanisms of information storage in visual short-term memory. Vision research. 2016;128:53–67. Epub 20161004. doi: 10.1016/j.visres.2016.09.010. PubMed PMID: 27668990; PMCID: PMC5079778.

31. Christophel TB, Klink PC, Spitzer B, Roelfsema PR, Haynes JD. The Distributed Nature of Working Memory. Trends in cognitive sciences. 2017;21(2):111–24. Epub 20170104. doi: 10.1016/j.tics.2016.12.007. PubMed PMID: 28063661.

32. Gresch D, Boettcher SEP, Gohil C, van Ede F, Nobre AC. Neural dynamics of shifting attention between perception and working-memory contents. Proceedings of the National Academy of Sciences of the United States of America. 2024;121(47):e2406061121. Epub 20241113. doi: 10.1073/pnas.2406061121. PubMed PMID: 39536078.

33. Jones HM, Diaz GK, Ngiam WXQ, Awh E. Electroencephalogram Decoding Reveals Distinct Processes for Directing Spatial Attention and Encoding Into Working Memory. Psychological science. 2024;35(10):1108–38. Epub 20240819. doi: 10.1177/09567976241263002. PubMed PMID: 39159181.

34. Berggren N, Eimer M. Does Contralateral Delay Activity Reflect Working Memory Storage or the Current Focus of Spatial Attention within Visual Working Memory? Journal of cognitive neuroscience. 2016;28(12):2003–20. Epub 20160726. doi: 10.1162/jocn_a_01019. PubMed PMID: 27458749.

35. Hakim N, Adam KCS, Gunseli E, Awh E, Vogel EK. Dissecting the Neural Focus of Attention Reveals Distinct Processes for Spatial Attention and Object-Based Storage in Visual Working Memory. Psychological science. 2019;30(4):526–40. Epub 20190228. doi: 10.1177/0956797619830384. PubMed PMID: 30817220; PMCID: PMC6472178.

36. Panichello MF, Buschman TJ. Shared mechanisms underlie the control of working memory and attention. Nature. 2021;592(7855):601–5. Epub 20210331. doi: 10.1038/s41586-021-03390-w. PubMed PMID: 33790467; PMCID: PMC8223505.

37. Mendoza-Halliday D, Xu H, Azevedo FAC, Desimone R. Dissociable neuronal substrates of visual feature attention and working memory. Neuron. 2024;112(5):850–63 e6. Epub 20240115. doi: 10.1016/j.neuron.2023.12.007. PubMed PMID: 38228138; PMCID: PMC10939754.

38. Ester EF, Sprague TC, Serences JT. Parietal and Frontal Cortex Encode Stimulus-Specific Mnemonic Representations during Visual Working Memory. Neuron. 2015;87(4):893–905. Epub 20150806. doi: 10.1016/j.neuron.2015.07.013. PubMed PMID: 26257053; PMCID: PMC4545683.

39. Rademaker RL, Chunharas C, Serences JT. Coexisting representations of sensory and mnemonic information in human visual cortex. Nature neuroscience. 2019;22(8):1336–44. Epub 20190701. doi: 10.1038/s41593-019-0428-x. PubMed PMID: 31263205; PMCID: PMC6857532.

40. Fiebelkorn IC, Kastner S. A Rhythmic Theory of Attention. Trends in cognitive sciences. 2019;23(2):87–101. doi: 10.1016/j.tics.2018.11.009. PubMed PMID: 30591373.

41. Benedetto A, Morrone MC, Tomassini A. The common rhythm of action and perception. Journal of cognitive neuroscience. 2019;32(2):187–200. doi: 10.1162/jocn_a_01436.

42. Fiebelkorn IC, Saalmann YB, Kastner S. Rhythmic sampling within and between objects despite sustained attention at a cued location. Current biology : CB. 2013;23(24):2553–8. doi: 10.1016/j.cub.2013.10.063. PubMed PMID: 24316204; PMCID: 3870032.

43. Helfrich RF, Fiebelkorn IC, Szczepanski SM, Lin JJ, Parvizi J, Knight RT, Kastner S. Neural mechanisms of sustained attention are rhythmic. Neuron. 2018;99(4):829–41. doi: 10.1016/j.neuron.2018.07.032.

44. Fiebelkorn IC, Pinsk MA, Kastner S. A dynamic interplay within the frontoparietal network underlies rhythmic spatial attention. Neuron. 2018;99(4):842–53. doi: 10.1016/j.neuron.2018.07.038.

45. Landau AN, Fries P. Attention samples stimuli rhythmically. Current biology : CB. 2012;22(11):1000–4. Epub 2012/05/29. doi: 10.1016/j.cub.2012.03.054. PubMed PMID: 22633805.

46. Landau AN, Schreyer HM, van Pelt S, Fries P. Distributed Attention Is Implemented through Theta-Rhythmic Gamma Modulation. Current biology : CB. 2015;25(17):2332–7. doi: 10.1016/j.cub.2015.07.048. PubMed PMID: 26279231.

47. Dugue L, Marque P, VanRullen R. Theta oscillations modulate attentional search performance periodically. Journal of cognitive neuroscience. 2015;27(5):945–58. doi: 10.1162/jocn_a_00755. PubMed PMID: 25390199.

48. Dugue L, Roberts M, Carrasco M. Attention Reorients Periodically. Current biology : CB. 2016;26(12):1595–601. doi: 10.1016/j.cub.2016.04.046. PubMed PMID: 27265395; PMCID: 4935543.

49. VanRullen R, Carlson T, Cavanagh P. The blinking spotlight of attention. Proceedings of the National Academy of Sciences of the United States of America. 2007;104(49):19204–9. doi: 10.1073/pnas.0707316104. PubMed PMID: 18042716; PMCID: 2148268.

50. Busch NA, VanRullen R. Spontaneous EEG oscillations reveal periodic sampling of visual attention. Proceedings of the National Academy of Sciences of the United States of America. 2010;107(37):16048–53. Epub 2010/09/02. doi: 10.1073/pnas.1004801107. PubMed PMID: 20805482; PMCID: 2941320.

51. Abdalaziz M, Redding ZV, Fiebelkorn IC. Rhythmic temporal coordination of neural activity prevents representational conflict during working memory. Current biology : CB. 2023;33(9):1855–63 e3. Epub 20230425. doi: 10.1016/j.cub.2023.03.088. PubMed PMID: 37100058.

52. Siegel M, Warden MR, Miller EK. Phase-dependent neuronal coding of objects in short-term memory. Proceedings of the National Academy of Sciences of the United States of America. 2009;106(50):21341–6. doi: 10.1073/pnas.0908193106. PubMed PMID: 19926847; PMCID: 2779828.

53. Bahramisharif A, Jensen O, Jacobs J, Lisman J. Serial representation of items during working memory maintenance at letter-selective cortical sites. PLoS biology. 2018;16(8):e2003805. Epub 20180815. doi: 10.1371/journal.pbio.2003805. PubMed PMID: 30110320; PMCID: PMC6093599.

54. Peters B, Kaiser J, Rahm B, Bledowski C. Object-based attention prioritizes working memory contents at a theta rhythm. Journal of experimental psychology General. 2021;150(6):1250–6. Epub 20201119. doi: 10.1037/xge0000994. PubMed PMID: 33211526.

55. Kaminski J, Brzezicka A, Mamelak AN, Rutishauser U. Combined Phase-Rate Coding by Persistently Active Neurons as a Mechanism for Maintaining Multiple Items in Working Memory in Humans. Neuron. 2020;106(2):256–64 e3. Epub 20200220. doi: 10.1016/j.neuron.2020.01.032. PubMed PMID: 32084331; PMCID: PMC7217299.

56. Chota S, Leto C, van Zantwijk L, Van der Stigchel S. Attention rhythmically samples multi-feature objects in working memory. Scientific reports. 2022;12(1):14703. Epub 20220829. doi: 10.1038/s41598-022-18819-z. PubMed PMID: 36038570; PMCID: PMC9424255.

57. Pomper U, Ansorge U. Theta-Rhythmic Oscillation of Working Memory Performance. Psychological science. 2021;32(11):1801–10. Epub 20210930. doi: 10.1177/09567976211013045. PubMed PMID: 34592108.

58. Hogendoorn H. Voluntary Saccadic Eye Movements Ride the Attentional Rhythm. Journal of cognitive neuroscience. 2016;28(10):1625–35. doi: 10.1162/jocn_a_00986. PubMed PMID: 27243615.

59. Souza AS, Oberauer K. In search of the focus of attention in working memory: 13 years of the retro-cue effect. Attention, perception & psychophysics. 2016;78(7):1839–60. doi: 10.3758/s13414-016-1108-5. PubMed PMID: 27098647.

60. Maris E, Oostenveld R. Nonparametric statistical testing of EEG- and MEG-data. Journal of neuroscience methods. 2007;164(1):177–90. Epub 20070410. doi: 10.1016/j.jneumeth.2007.03.024. PubMed PMID: 17517438.

61. Oostenveld R, Fries P, Maris E, Schoffelen JM. FieldTrip: Open source software for advanced analysis of MEG, EEG, and invasive electrophysiological data. Computational intelligence and neuroscience. 2011;2011:156869. doi: 10.1155/2011/156869. PubMed PMID: 21253357; PMCID: 3021840.

62. Perrin F, Pernier J, Bertrand O, Giard MH, Echallier JF. Mapping of scalp potentials by surface spline interpolation. Electroencephalography and clinical neurophysiology. 1987;66(1):75–81. doi: 10.1016/0013-4694(87)90141-6. PubMed PMID: 2431869.

63. Fiebelkorn IC, Snyder AC, Mercier MR, Butler JS, Molholm S, Foxe JJ. Cortical cross-frequency coupling predicts perceptual outcomes. NeuroImage. 2013;69:126–37. doi: 10.1016/j.neuroimage.2012.11.021. PubMed PMID: 23186917; PMCID: 3872821.

64. Fiebelkorn IC, Pinsk MA, Kastner S. The mediodorsal pulvinar coordinates the macaque fronto-parietal network during rhythmic spatial attention. Nature communications. 2019;10(1):215. doi: 10.1038/s41467-018-08151-4. PubMed PMID: 30644391.

65. Bosman CA, Womelsdorf T, Desimone R, Fries P. A microsaccadic rhythm modulates gamma-band synchronization and behavior. The Journal of neuroscience : the official journal of the Society for Neuroscience. 2009;29(30):9471–80. doi: 10.1523/JNEUROSCI.1193-09.2009. PubMed PMID: 19641110.

66. Hillyard SA, Anllo-Vento L. Event-related brain potentials in the study of visual selective attention. Proceedings of the National Academy of Sciences of the United States of America. 1998;95(3):781–7. doi: 10.1073/pnas.95.3.781. PubMed PMID: 9448241; PMCID: PMC33798.

67. O’Connell MN, Falchier A, McGinnis T, Schroeder CE, Lakatos P. Dual mechanism of neuronal ensemble inhibition in primary auditory cortex. Neuron. 2011;69(4):805–17. Epub 2011/02/23. doi: 10.1016/j.neuron.2011.01.012. PubMed PMID: 21338888; PMCID: 3052772.

68. O’Connell RG, Dockree PM, Kelly SP. A supramodal accumulation-to-bound signal that determines perceptual decisions in humans. Nature neuroscience. 2012;15(12):1729–35. Epub 20121028. doi: 10.1038/nn.3248. PubMed PMID: 23103963.

69. VanRullen R. How to Evaluate Phase Differences between Trial Groups in Ongoing Electrophysiological Signals. Frontiers in neuroscience. 2016;10:426. Epub 20160914. doi: 10.3389/fnins.2016.00426. PubMed PMID: 27683543; PMCID: PMC5021700.

70. Senoussi M, Moreland JC, Busch NA, Dugue L. Attention explores space periodically at the theta frequency. Journal of vision. 2019;19(5):22. doi: 10.1167/19.5.22. PubMed PMID: 31121012.

71. Gresch D, Boettcher SEP, van Ede F, Nobre AC. Shifting attention between perception and working memory. Cognition. 2024;245:105731. Epub 20240125. doi: 10.1016/j.cognition.2024.105731. PubMed PMID: 38278040.

72. Foxe JJ, Snyder AC. The Role of Alpha-Band Brain Oscillations as a Sensory Suppression Mechanism during Selective Attention. Frontiers in psychology. 2011;2:154. doi: 10.3389/fpsyg.2011.00154. PubMed PMID: 21779269; PMCID: 3132683.

73. Gresch D, Boettcher SEP, van Ede F, Nobre AC. Shielding working-memory representations from temporally predictable external interference. Cognition. 2021;217:104915. Epub 20210929. doi: 10.1016/j.cognition.2021.104915. PubMed PMID: 34600356; PMCID: PMC8543071.

74. Verschooren S, Pourtois G, Egner T. More efficient shielding for internal than external attention? Evidence from asymmetrical switch costs. Journal of experimental psychology Human perception and performance. 2020;46(9):912–25. Epub 20200507. doi: 10.1037/xhp0000758. PubMed PMID: 32378935.

